# The cell type composition of the adult mouse brain revealed by single cell and spatial genomics

**DOI:** 10.1101/2023.03.06.531307

**Authors:** Jonah Langlieb, Nina S. Sachdev, Karol S. Balderrama, Naeem M. Nadaf, Mukund Raj, Evan Murray, James T. Webber, Charles Vanderburg, Vahid Gazestani, Daniel Tward, Chris Mezias, Xu Li, Dylan M. Cable, Tabitha Norton, Partha Mitra, Fei Chen, Evan Z. Macosko

## Abstract

The function of the mammalian brain relies upon the specification and spatial positioning of diversely specialized cell types. Yet, the molecular identities of the cell types, and their positions within individual anatomical structures, remain incompletely known. To construct a comprehensive atlas of cell types in each brain structure, we paired high-throughput single-nucleus RNA-seq with Slide-seq–a recently developed spatial transcriptomics method with near-cellular resolution–across the entire mouse brain. Integration of these datasets revealed the cell type composition of each neuroanatomical structure. Cell type diversity was found to be remarkably high in the midbrain, hindbrain, and hypothalamus, with most clusters requiring a combination of at least three discrete gene expression markers to uniquely define them. Using these data, we developed a framework for genetically accessing each cell type, comprehensively characterized neuropeptide and neurotransmitter signaling, elucidated region-specific specializations in activity-regulated gene expression, and ascertained the heritability enrichment of neurological and psychiatric phenotypes. These data, available as an online resource (BrainCellData.org) should find diverse applications across neuroscience, including the construction of new genetic tools, and the prioritization of specific cell types and circuits in the study of brain diseases.

---

The mammalian brain is composed of a remarkably diverse array of cell types that display high degrees of molecular, anatomical and physiological specialization. Although the precise number of distinct cell types present in the brain is unknown, the number is presumed to be in the thousands^1, 2^. These cell types are the building blocks of hundreds of discrete neuroanatomical structures^3^, each of which plays a distinct role in brain function. Advances in the throughput of single-cell RNA sequencing (scRNAseq) technology has enabled the generation of cell type inventories in many individual brain regions^4–13^, as well as the construction of broader atlases that coarsely cover the nervous system^14, 15^. Furthermore, application of new spatial transcriptomics techniques to the brain has begun to illuminate the spatial organization of brain cell types^10, 16–18^. However, a full inventory of cell types across the brain, with their cell bodies localized to specific neuroanatomical structures, does not yet exist.

## Transcriptional diversity and cell type representation across neuroanatomical structures

To comprehensively sample cell types across the brain, we employed a recently developed pipeline for high-throughput single-nucleus RNA-seq (snRNA-seq) that has high transcript capture efficiency and nuclei recovery efficiency, as well as consistent performance across diverse brain regions^6, 7^. We dissected and isolated single nuclei from 92 discrete anatomical locations, derived from 55 individual mice (Fig. 1a, Methods, Supplemental Table 1). Across all 92 dissectates, after all quality control steps (Methods), we recovered a total of 4,388,420 nuclei profiles with a median transcript capture of 4,884 unique molecular identifiers (UMIs) per profile (Extended Data Fig. 1a-d). We sampled nearly equal numbers of profiles from male and female donors, with minimal batch effects across mice, such that replicates of individual dissectates contributed to each cluster (Extended Data Fig. 1e). To discover cell types, we developed a simplified iterative clustering strategy in which the cells were repeatedly clustered on distinctions amongst a small set of highly variable genes until clusters no longer could be distinguished by at least 3 discrete markers (Methods). Our clustering algorithm largely recapitulated published results of the motor cortex^4^ and cerebellum^7^ (Extended Data Fig. 1f), and was scalable to support the computational analysis of millions of cells (Methods). In total, after quality control and cluster annotation (Methods), we identified 4,998 discrete clusters, the great majority of which (97%) were neuronal (Fig. 1a, Extended Data Fig. 1g), consistent with prior large-scale surveys of brain cell types^14, 15^. Across the brain, we estimate that our sampling depth reached an estimated 90% saturation of cell type discovery (Methods, Extended Data Fig. 1h).

**Figure 1.**
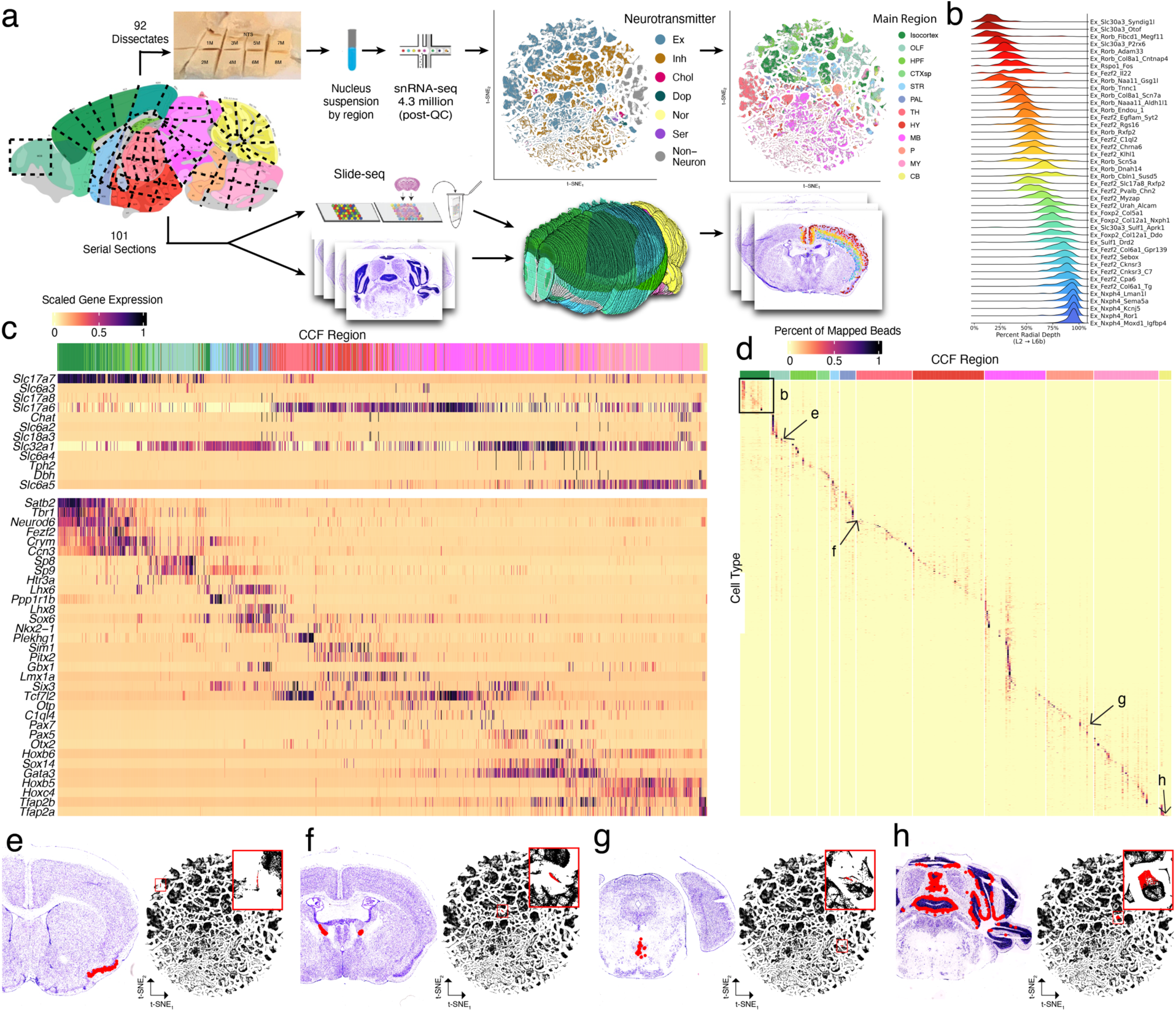
Spatially mapping cell types using whole-brain snRNA-seq and Slide-seq datasets. **a**, Schematic of the experimental and computational workflows for both whole-brain snRNA-seq sampling (top arrows) and Slide-seq sampling (bottom arrows). Top, t-SNE representations of gene expression relationships amongst 1.2 million spatially mapped snRNA-seq profiles (downsampled from 4.3 million) colored by neurotransmitter identity (left) and most common spatially mapped main region (right). **b**, Ridge plot depicting the spatial distributions of excitatory cortical cell types along the laminar depth of cortex in the Slide-seq dataset. **c**, Heatmap depicting expression of main neurotransmitter genes (top) and canonical neuronal cell type markers (bottom) across all 1,260 spatially mapped neuronal clusters. **d**, Heat map showing the spatial distributions of each spatially mapped cluster (rows) within each DeepCCF structure (for complete list, see Supplemental Table 2). Example mapped cell types in other panels are labeled on the heat map. **e-h**, Example mappings of neuronal cell types throughout the brain plotted in the CCF-aligned Slide-seq data (left) and in t-SNE space (with insets).

To determine the spatial distributions of these cell types, we next performed Slide-seq^19, 20^ on serial coronal sections of one hemisphere of an adult female mouse brain (Methods), spaced ∼100 μm, matching the resolution of commonly used neuroanatomical atlases^21, 22^. Slide-seq detects the expression of genes on 10 μm beads across the transcriptome within a fresh-frozen tissue section, providing near-cellular resolution data. In total, we sequenced 101 arrays, spanning the entire anterior-posterior axis of the brain. We aligned the sequencing-generated Slide-seq images to adjacent Nissl sections, which were themselves aligned to the Allen Common Coordinate Framework^3^ (CCF Framework) (Methods, Extended Data Fig. 2a), enabling us to assign a neuroanatomical structure to each bead (Fig. 1a). To confirm the accuracy of our alignment, we plotted expression of three highly region-specific markers across our CCF-defined regions, and quantified the distance of each expressing bead from the expected CCF region (Extended Data Fig. 2b). From this analysis, we estimate our alignment error to be in the range of 22 – 94 μm (Extended Data Fig. 2c).

To localize cell types to brain structures, we computationally decomposed individual Slide-seq beads into combinations of snRNA-seq-defined cluster signatures using Robust Decomposition of Cell Type Mixtures (RCTD)^23^. To handle the enormous cellular complexity of these regions, we implemented RCTD in a highly parallelized computational environment^24^, and developed a confidence score that more accurately distinguishes among groups of highly similar cell type definitions (Methods). In total, we mapped 1,931 snRNA-seq-defined clusters (Methods) to greater than 1.7 million beads within the Slide-seq dataset. We computed the cortical layer depth of a set of 42 isocortical excitatory neuronal types and found the mappings had the expected highly-regionalized radial depth^5^ (Fig. 1b) when ordered by their best integrated match with a previous cortical atlas^5^, suggesting faithful projection of cell type signatures into spatial coordinates.

We next evaluated the consistency and comprehensiveness of our cluster set by plotting known markers of cell-type identity across the clusters (Methods). We assessed the consistency of expression of these known markers (mostly transcription factors) with the expected localizations of cell types across twelve main brain regions defined in the Allen Brain Atlas (Fig. 1c): isocortex, the olfactory areas (OLF), hippocampal formation (HPF), striatum (STR), pallidum (PAL), hypothalamus (HY), thalamus (TH), midbrain (MB), pons (P), medulla (MY), and cerebellum (CB). Amongst our neuronal clusters, we identified cortical, amygdalar, olfactory, and hippocampal excitatory projection neurons (*Tbr1*, *Neurod6*, and *Satb2*); telencephalic interneurons (*Sp8*, *Sp9*, and *Htr3a*); spiny projection neurons of the striatum and adjacent pallidal structures (*Ppp1r1b*); hypothalamic neurons (*Nkx2-1*, *Sim1, Lhx6,* and *Lhx8*); principal neurons of the thalamus (*Tcf7l2*, *Six3*, and *Plekhg1*); neurons of the brainstem that populate mostly midbrain and pontine structures (*Otx2*, *Gata3*, *Pax5, Pax7*, and *Sox14*); neurons expressing *Hox* homeobox genes that are primarily in the rhombencepthalon; and cerebellar neurons expressing *Tfap2a* and *Tfap2b*. Neurons also specialize in the specific neurotransmitters they release. We detected discrete populations of gluatmatergic (*Slc17a6*, *Slc17a7*, and *Slc17a8*), GABAergic (*Slc32a1*), glycinergic (*Slc6a5*), cholinergic (*Chat* and *Slc18a3*), serotonergic (*Slc6a4* and *Tph2*), dopaminergic (*Slc6a3*), and noradrenergic (*Slc6a2* and *Dbh*) cell types, distributed in the expected regions. Together, these results indicate that our systematic sampling covered the expected molecular diversity of neurons across the mouse brain.

Many glial populations were distributed across large neuroanatomical boundaries (telencephalon, mesencephalon, and rhombencephalon), indicating that, relative to neurons, regional gene expression differences amongst glial populations were small (Extended Data Fig. 2d). Indeed, neuronal populations mapped to much more specific and related structures (Fig. 1b,d-h), reflecting the strong regional specificity of neuronal specializations. We assessed the distribution of neuronal cell types within a more granular set of 256 CCF (termed “DeepCCF”) structures (Supplemental Table 2). Most cell types showed highly refined regional localization: 60% of mapped clusters were confidently mapped (Methods) in 3 or fewer DeepCCF regions, reflecting the extent to which neuroanatomical nuclei are individually composed of locally diversified cell types.

## Variation in neuronal diversity across neuroanatomical structures

Our initial results revealed surprisingly large numbers of cell types distributed across the main brain regions. To explore cellular diversity at a finer neuroanatomical scale, we tallied the number of cell types confidently mapping to each DeepCCF structure, computing the number of types needed to occupy 95% of all mapped beads localized within that DeepCCF structure (Methods). Within the main 12 brain regions, we found the largest diversity of cell types in the midbrain, followed by hypothalamus, pons, and medulla (Fig. 2a). Within the more fine-grained DeepCCF structures, we found particularly high cell type diversity within the periaqueductal gray matter (PAG) and midbrain reticular nucleus (MRN) of the midbrain. Regions of high diversity in other major brain areas included the parvicellular reticular nucleus of the medulla (PARN), the pontine reticular nucleus (PRNr), the lateral hypothalamic area (LHA), and the bed nucleus of the stria terminalis (BST), consistent with our prior analysis of this area^6^. Although cell types were often highly focal within DeepCCF structures (Fig. 1b, d-h), some cell types also crossed DeepCCF boundaries. To visualize cellular compositional relationships amongst brain regions in greater detail, we built a force-directed graph in which the edges between DeepCCF regions were weighted to represent the number of clusters that jointly mapped in those regions (Methods, Fig. 2b). Cell types largely were restricted to each major brain area, but showed greater mixing between pons and medulla than other regions, indicating more mixing of cell types specifically within those structures (Extended Data Fig. 3a).

**Figure 2.**
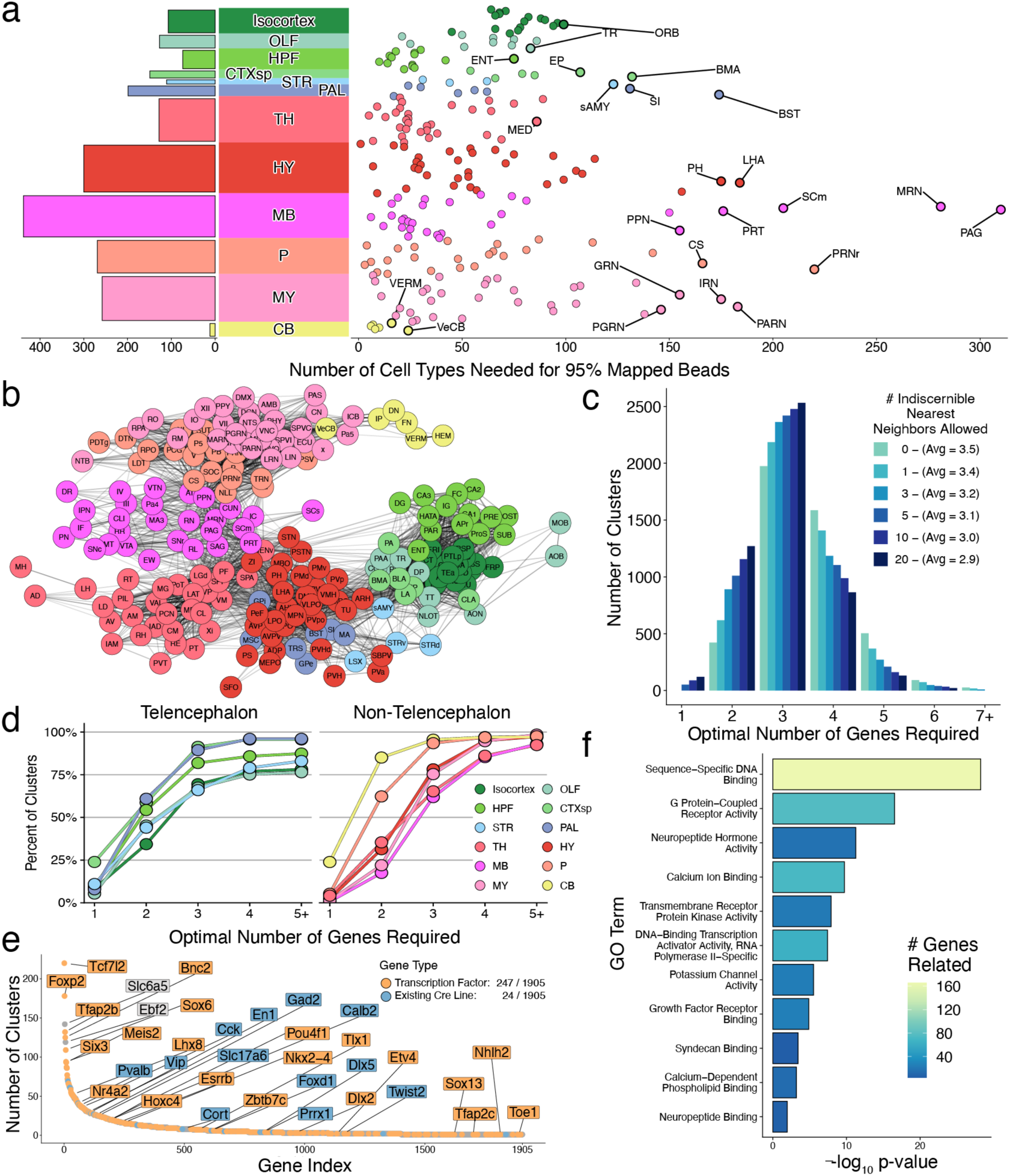
Cell type diversity across regions of the mouse brain. **a**, Cumulative number of cell types needed to reach 95% of mapped beads in each DeepCCF region (right) and summarized across individual main regions (left). The DeepCCF regions with the largest values are labeled. **b,** Force-directed graph showing cell type sharing relationships amongst DeepCCF regions. Edges are weighted by the Jaccard overlap between each region (Methods). **c,** Histogram of the minimum number of genes required to uniquely define each cell type across the nervous system. The algorithm was repeated, tolerating genes to be the same amongst different numbers of nearest cluster neighbors. **d**, Cumulative distribution plot of minimum number of genes required to distinguish each cell type within each major brain region, grouped by telencephalic regions (left), and non-telencephalic regions (right). **e**, Optimally-small collated gene set needed to cover a minimal gene list from each cell type, ranked by how many cell types each gene reaches, colored by transcription factor identity (orange) or if it is a currently available Cre line (blue). **f**, Barplot quantifying the significantly enriched GO terms in the minimum-sized collated gene list after hierarchical reduction (Methods), colored by the absolute number of genes related.

Circuit-level analyses of the mouse brain have relied upon the availability of genetically delivered molecular tools to excite, inhibit, and record from individual neuronal populations. These tools have historically been delivered to specific subpopulations of neurons through the use of recombinase-based systems, but more recently, RNA editing-based strategies have been developed to enable translation of transgenes only in the presence of specific endogenous mRNA transcripts^25–27^. Both strategies require nominating small numbers of high-value marker genes that can optimally distinguish amongst many distinct clusters. To identify the minimum number of genes needed to combinatorially define each cell type in our snRNA-seq dataset, we framed the question as a set cover problem^28^ (Methods), which can be solved to optimality using mixed integer linear programming techniques^29, 30^. For a great majority of cell types (93%), our algorithm was able to recover a minimally sized set of defining genes (all combinations are available in Supplemental Table 3).

Examination of the overall results revealed that across the 4,990 clusters used, the median minimally sized gene set was 3. Depending on the tolerance we set for the number of neighbors that co-express the same selected gene markers, between 13 and 89 cell types could be uniquely defined by a single gene, while between 422 and 1,156 cell types could be uniquely defined by two genes. When we performed the analysis on each of the 12 major brain regions separately, more than 70% of cell types could be uniquely defined, within that area, by up to 3 genes (Fig. 2c, Extended Data Fig. 3b). However, there was considerable variation in these minimal gene sets when stratifying by brain region: more than 45% of cell types within telencephalic brain areas could be uniquely defined by up to two genes, while in the midbrain and medulla only 17% and 22% could be, respectively (Fig. 2d). These results suggest that the molecular diversity of brainstem areas is especially combinatorial, and motivates the development of molecular strategies that can provide intersectional expression gating in order to properly isolate them and characterize their unique functions.

We also discovered that most cell types had more than one distinct minimally sized gene list. We leveraged this redundancy to determine the smallest set of genes that would encompass at least one gene list from each cell type (Methods). We identified a set of 1,905 genes that covered each cell type (Fig. 2e). This minimum gene set included many genes with existing Cre lines^31^, and was highly enriched for transcription factors (OR=2.54, p < 0.001), G-protein coupled receptors (OR=1.83, p < 0.001), and neuropeptides (OR=5.76, p < 0.001) (Fig. 2f, Extended Data Fig. 3c), gene families that have been historically used to define cell types in the brain.

## Principles of neurotransmission and neuropeptide usage

Neurons communicate with each other across synapses, through the release of different small molecules and peptides. We asked in which regions, and in which combinations, neurotransmitters are used across the cell types of the brain. Since the production and usage of these neurotransmitters at synapses requires different sets of gene products, we leveraged our snRNA-seq data to assign neurotransmitter identities to each cell type (Methods).

Overall, amongst the neuronal snRNA-seq clusters, cell type diversity was well balanced between excitatory and inhibitory cell types (2,420 excitatory, 2,246 inhibitory), and co-transmission of glutamate with an inhibitory neurotransmitter (GABA or glycine) was relatively rare (1.1% of all neuronal clusters, Fig. 3a). Most co-releasing populations (35 out of 54) expressed the glutamate transporter *Slc17a8* (VGLUT3) and derived from a wide range of lineages, populating regions across the telencephalon, midbrain, and hindbrain. Amongst neuron types releasing neuromodulators, we found the cholinergic neurons were more diverse (102 clusters) than serotonergic and dopaminergic types (25 and 13 clusters respectively), and were distributed much more widely across the nervous system (Extended Data Fig. 4a,b).

**Figure 3.**
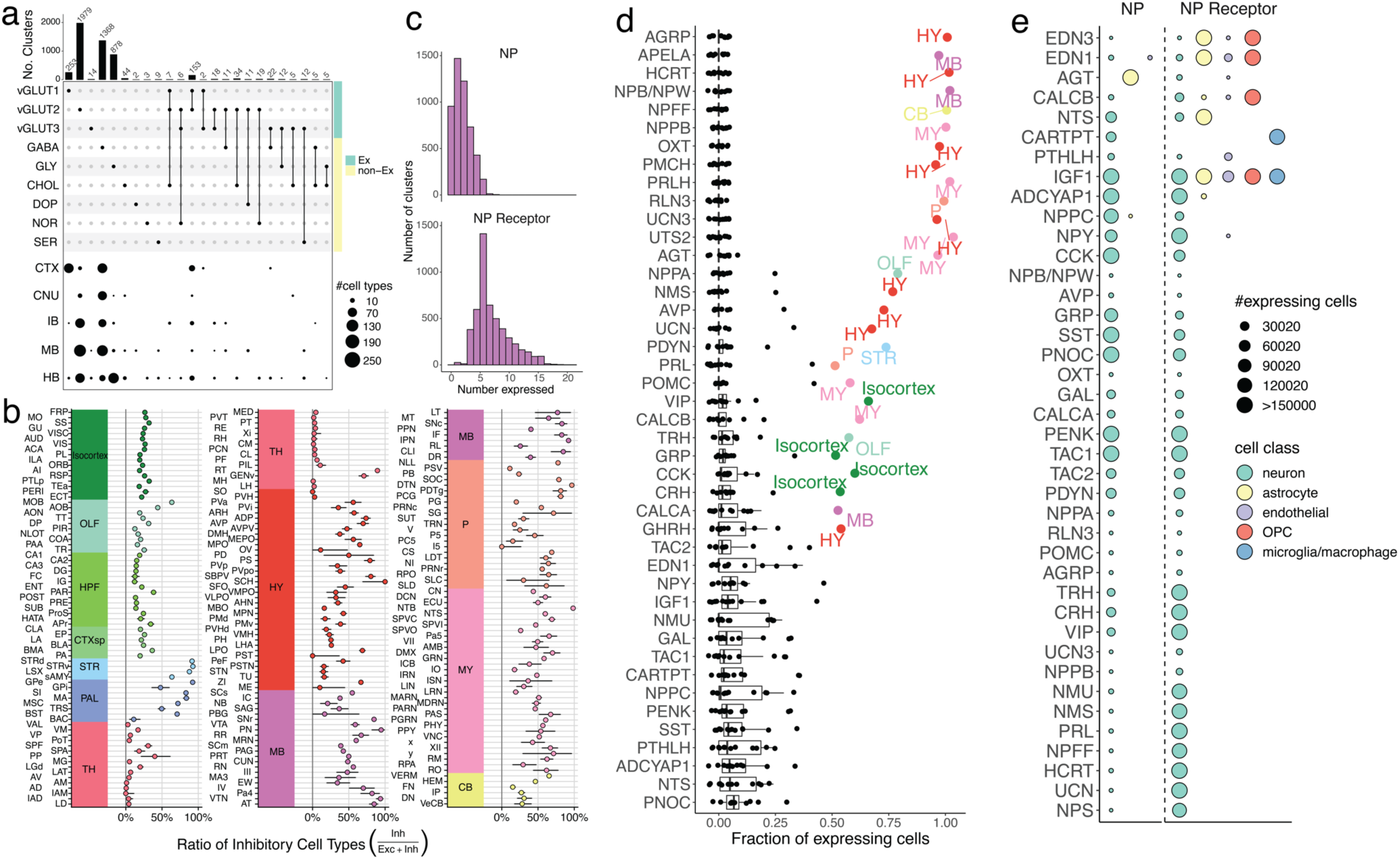
Neurotransmission and neuropeptide (NP) usage across regions of the mouse brain. **a**, Top, upset plot of the frequency of neurotransmitter usage by individual snRNA-seq-defined cell types. Bottom, dot plot depicting the spatial distribution of cell types in each of the neurotransmitter groups across major brain areas. **b**, Point estimates of the fraction of each DeepCCF region composed of mapped inhibitory cell types. Whiskers denote the exact 95% confidence interval of the corresponding binomial distribution. **c**, Histograms denoting the number of distinct NPs (top) and NP receptors (bottom) expressed in each snRNA-seq-defined neuronal cell type. **d**, Fraction of all cells expressing each NP (y-axis) in each of the 12 main brain areas. Regions accounting for more than 50% of total expression of that NP are colored and labeled. **e**, Dot plot depicting the number of cells expressing each NP (left of dotted line) and NP receptor (right of dotted line), within each major cell class.

Although the brain-wide cellular composition was balanced between inhibitory and excitatory types, individual brain regions are known to be composed of more skewed compositions of excitatory or inhibitory neurons. To characterize neurotransmission balance comprehensively in all structures, we quantified the excitatory-to-inhibitory (E-I) balance of each DeepCCF region by comparing the ratio of the number of beads mapping to excitatory cell types, to those mapping to inhibitory cell types (Methods). The computed E-I balances recovered the expected broad patterns, including the dominance of excitatory cells in thalamic nuclei, and the lack of excitatory populations within the striatum (Fig. 3b). Furthermore, more subtle distinctions could also be appreciated, such as the higher inhibitory proportion in certain thalamic nuclei known to contain interneurons (e.g. LGd). Within the telencephalon, regions were more commonly skewed towards a predominantly excitatory (e.g. cortical regions), or predominantly inhibitory (e.g. striatum) composition. In addition, regions with high E-I imbalance were more likely to be predominantly excitatory, while predominantly inhibitory regions were less common, being largely restricted to the striatum, the thalamic reticular nucleus, and a few brainstem nuclei.

Neuropeptides (NPs) exert varied and complex neuromodulatory effects on circuits through downstream G-protein coupled receptors (GPCRs). Neuropeptides are also often co-released with other neurotransmitters to directly modulate synaptic activity. We utilized our spatially mapped cell type inventory to characterize the basic rules and principles by which NPs are used throughout the brain. We curated a set of 61 genes that produce at least one NP with a known downstream GPCR (Supplemental Table 4), and quantified the number of NP-expressing, and NP-sensing, cell types. Amongst our 4,998 cell types, 80.9% expressed at least one neuropeptide, underscoring the ubiquity of NP signaling in the mammalian central nervous system (Fig. 3c). Neuropeptide sensing was even more ubiquitous: 91.6% of cell types expressed receptors for more than three NPs. Historically, NP signaling has been particularly strongly associated with the hypothalamus, where many of the NPs were originally biochemically discovered^32^. However, our analyses did not find that, overall, hypothalamic neurons were any more likely to release neuropeptides, compared with neurons in other brain areas (Extended Data Fig. 4c). Rather, the hypothalamus, as well as the pallidum and midbrain, were more likely to release a subset of neuropeptides–like oxytocin or vasopressin– that are highly selectively expressed, while other brain regions expressed NPs that were more ubiquitous throughout the nervous system (Fig. 3d).

Nearly all neuropeptides were both expressed and sensed by neuronal cell types (Fig. 3e). However, we identified two likely examples of NP signaling between neurons and glia. The expression of *Cartpt* was detected in 232 neuronal populations, distributed in hypothalamic and midbrain regions, while its receptor *Gpr160*^33^ was highly restricted to microglia and macrophage populations. Interestingly, *Gpr160* induction was observed to be within microglia in a recent study of spinal cord nerve injury^33^. Conversely, the expression of the angiotensin-encoding gene *Agt* was found to be primarily in astrocytes found in non-telencephalic regions (Fig. 3e, Extended Data Fig. 4d), while its receptors *Agtr1a* and *Agtr2* were enriched in non-telencephalic neurons. Astrocyte-neuron signaling through angiotensin could play important homeostatic roles particularly in the midbrain, where DA neurons vulnerable to neurodegeneration in Parkinson’s disease (PD) were recently identified to selectively express *Agtr1a*^34^, and inhibition of the angiotensin receptor has been shown to be neuroprotective in PD animal models^35^ and in clinical cohorts^36^.

## Activity-dependent gene enrichment across cell types and regions

Neuronal cells, in response to an increase in action potential firing, induce the expression of hundreds of activity-regulated genes (ARGs)^37^. The prototypical ARG is *Fos*, which is induced within minutes of elevated activity, along with several highly correlated genes including *Junb* and *Egr1*, which are collectively referred to as immediate early genes (IEGs). These IEGs have been primarily discovered and studied in excitatory cortical or hippocampal cells. Our Slide-seq and snRNA-seq atlases provide two key advantages for assessing ARG heterogeneity across cell types. First, they are comprehensive in their coverage of the brain to enable broad comparative analysis. Second, they are performed on brain tissue that is frozen immediately after animal perfusion, eliminating any postmortem effects on ARG expression^38, 39^.

To characterize ARGs across neuronal types, we first partitioned our mapped clusters into 28 cell type groups defined by their Slide-seq mapped region, and their neurotransmitter identity (Methods). We then selected 407 candidate ARGs whose correlation with *Fos* was at least 0.3, met statistical significance (adjusted p-value < 0.05), and for which *Fos* was also above the 99.5% quantile of all correlations, in at least one cell type group (Methods). To ensure robustness, we validated that our candidate ARGs were similarly correlated with another canonical IEG, *Junb* (Extended Data Fig. 5a). To visualize which genes are consistently correlated across cell type groups, we constructed a bipartite graph, connecting each gene to cell type groups within which it is highly correlated with *Fos* (Methods, Fig. 4a). Examination of this graph revealed that the most connected genes–those that are most consistently and highly correlated with *Fos* across the brain–-included most canonical IEGs, such as *Egr1*, *Npas4*, *Arc*, *Junb*, *Btg2*, and *Nr4a1* (center of graph, Fig. 4a). We selected the eight most correlated of these genes to compare their relative activity across regions and cell types. Expression of these IEGs across each region in our Slide-seq dataset was highest in the isocortex, olfactory bulb, striatum, and amygdala, while regions of cerebellum and medulla showed the lowest average IEG expression (Fig. 4b, Extended Data Fig. 5b). Similarly, in our snRNA-seq clusters, IEG activity was noticeably higher in excitatory populations, particularly those in isocortex, olfactory areas, and hippocampal formation (Extended Data Fig. 5c).

**Figure 4.**
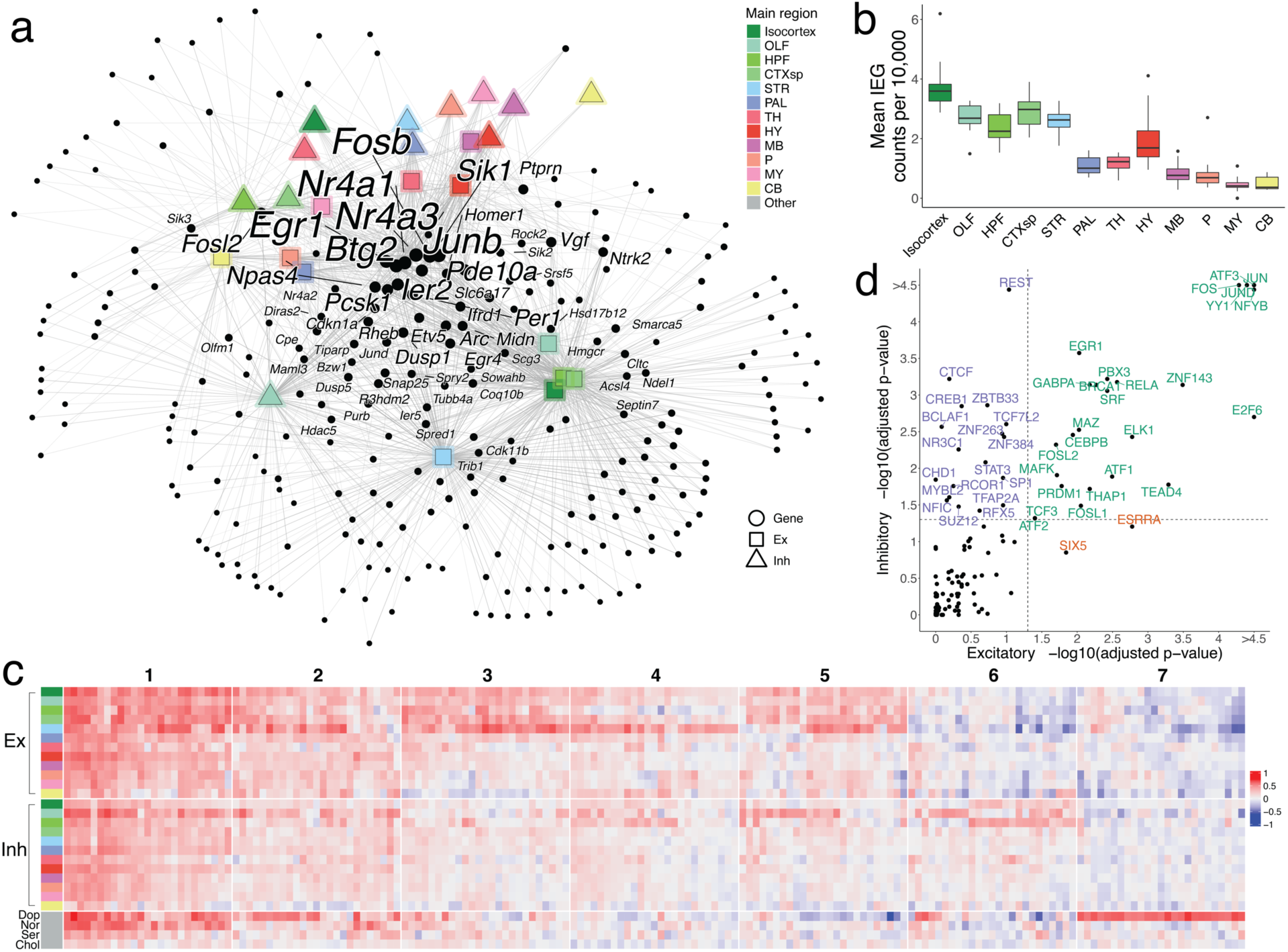
Patterns of activity-dependent gene expression across brain regions. **a**, Force-directed graph of a weighted bipartite network. Nodes comprise two disjoint sets: candidate ARGs (black dots) and sets of neuronal cell types of the same neurotransmitter type, localized to the same brain region (shapes colored by region). Edges are weighted based on the correlation coefficient between a gene node and a region node, and edges with weight < 1.3 are pruned. After pruning, nodes with a degree < 2 are also removed. The sizes of nodes and node labels correlate with the degree of the node, where nodes are labeled only if they share edges of weight ≥ 1.3 with at least six regions. **b**, Box plots quantifying mean core IEG Slide-seq counts per 10,000, colored by main brain regions. **c,** Downsampled heatmap of correlation coefficients between *Fos* and candidate ARGs (columns) across major regions of the brain (rows). Numbers at the top correspond to ARG cluster identities. **d,** Scatter plot quantifying transcription factor enrichment (p-value < 0.05, FDR-corrected) between excitatory and inhibitory populations. TFs are colored by their cell type enrichment specificity.

Our candidate ARG set also contained many genes connected to only a few of the major cell type groups, suggesting heterogeneity in the transcriptional programs of cell types in response to activity. To more deeply explore cell-type-specific ARGs, we hierarchically clustered our gene set into 7 clusters. Clusters 1-4 were the most universally correlated across cell types and regions (Fig. 4c, Extended Data Fig. 5d), and were highly enriched for known ARGs^40^ (Methods, Extended Data Fig. 5e), while clusters 5-7 were more cell-type-specific: cluster 5 was relatively specific for telencephalic excitatory neurons, cluster 6 was more specific for telencephalic inhibitory neurons, and cluster 7 was specific for dopaminergic neurons. Our inhibitory-specific cluster 6 included several genes previously reported as activity-regulated in cortical interneurons, such as *Crh* and *Cxcl14*^39^. Many of these genes are implicated in dendritic spine development and remodeling, such as *Tshr*^41^, *Shisa8*^42^, and *Sorcs2*^43^, indicating that synaptic plasticity may be a particularly prominent component of the activity-related response in telencephalic inhibitory cell types. To explore how the transcription of these gene sets may be differentially regulated across cell types, we compared the enrichment of transcription factor (TF) targets between genes highly correlated with *Fos* in either telencephalic excitatory or inhibitory cells (Methods). Amongst the 54 TFs with significant enrichment (p < 0.05, FDR-corrected; Fig. 4d), most (35 TFs) were jointly enriched in both inhibitory and excitatory populations, but inhibitory cells were selectively enriched for the targets of 15 TFs. These TFs included several well-known chromatin re-organizers, including CTCF, BCLAF1, and CHD1, suggesting an important role for epigenetic modification of inhibitory neurons in activity-dependent processes. Together, these analyses reveal how brain-wide, unbiased sampling of cell types can reveal not only the molecular markers defining these types, but also conserved, dynamic patterns of gene regulation that occur across cell type groups.

## Heritability enrichment of neurological and psychiatric traits

Over the past ten years, genome-wide association studies (GWAS) have uncovered risk loci associated with numerous neuropsychiatric traits. Identifying the cell types and brain regions in which these loci influence disease risk could catalyze new directions in understanding pathogenic mechanisms of many difficult-to-treat brain diseases. Because of their comprehensive coverage, our combined spatial and single-nucleus transcriptomics datasets provide a unique opportunity to investigate the relative enrichment of disease risk alleles across the entire mammalian nervous system. Several studies have integrated single cell and GWAS by aggregating cells from the same type and computing an enrichment statistic between the gene expression pattern of the cell type and the genes associated with risk by GWAS^9, 34, 44–46^. We used a recently described approach specifically developed for single-cell datasets^47^ (Methods) to evaluate the relative enrichment of loci from 16 neurological and psychiatric traits across our spatially localized cell types (Supplemental Table 5).

After multiple hypothesis correction testing (Methods), we identified a total of 154 cell types, across 11 traits, that met statistical significance (adjusted p-value < 0.05; Fig. 5a, Supplemental Table 6). The significance results were robust to using either pseudocells–aggregated collections of cellular neighborhoods that both reduce computational complexity and noise from statistical drop-out (Methods)–or individual cells (Extended Data Fig. 6a). For Alzheimer’s disease (AD), heritability enrichment was significant in macrophages and microglia, consistent with analyses of multiple prior datasets^34, 48, 49^. In autism spectrum disorder, two neuronal cell types showed statistically significant enrichment, distributed within the bed nucleus of the stria terminalis (BNST), an area with well-established roles in mediating social interactions, and the inferior colliculus, a midbrain structure involved in modulating auditory inputs, a common symptom of ASD patients. Educational attainment and major depressive disorder–two traits with known high polygenicity–showed enrichment across several regions (Extended Data Fig. 6b).

**Figure 5.**
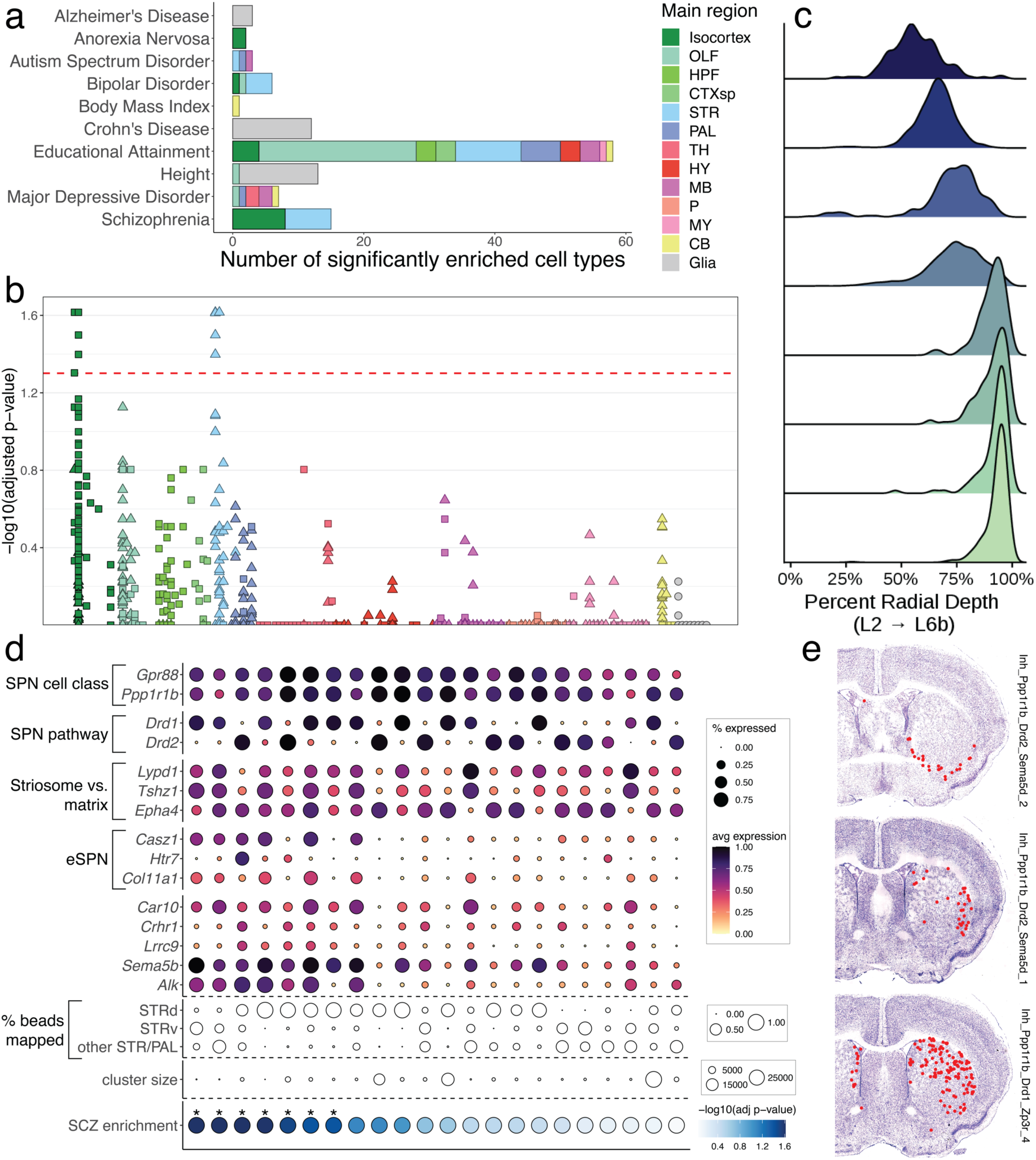
Heritability enrichment for traits studied by GWAS across brain cell types. **a**, Barplots quantifying significantly enriched (p-value < 0.05, FDR-corrected) cell types for each trait, in non-neurons (gray) and neurons (colored by main region). **b**, Adjusted −log10 p-value enrichment scores for each cell type, grouped and colored by their main regions, for schizophrenia. Squares and triangles denote excitatory and inhibitory clusters, respectively; glia are shown in gray on the far right of each plot. **c**, Ridge plots showing the layer distribution of each excitatory cortical cell type found to be enriched for schizophrenia heritability. **d**, Dot plot of expression of markers of striatal SPN subtype identity, grouped by category (overall cell class identity, pathway identity, matrix versus striosome, and eSPN identity). Five additional genes that are enriched in the schizophrenia-enriched SPN types are also shown. **e**, Representative sections showing the spatial localizations of three SPN cell types significantly enriched for schizophrenia heritability.

In schizophrenia (Fig. 5b) and bipolar disorder (Extended Data Fig. 6b), we observed enrichment signals within the excitatory neurons of the isocortex and the inhibitory neurons of the striatum, consistent both with the known shared heritability between these two disorders,^50, 51^ and prior enrichment studies performed on more limited collections of single cell datasets^45^. Importantly, although these two signals rose above our stringent threshold for multiple hypothesis testing correction, numerous other sub-threshold signals were present, suggesting that these cell type groups are not the only neuronal populations harboring enrichment for GWAS-associated genes. The significantly enriched excitatory populations were restricted to the lower layers (layers 5 and 6) of cortex (Fig. 5c), and expressed markers suggestive of intratelencephalic and layer 6b identities (Extended Data Fig. 6c). The enriched striatal neuron types all expressed the marker gene *Ppp1r1b*, identifying them as medium spiny neurons, the principal projection neurons (SPNs) of the dorsal and ventral striatum, which also populate several other pallidal structures. The SPNs can be subdivided by their projection pathway (indirect versus direct), their spatial localization^52^ (to the striatal matrix or striosome compartments), or more recently, by molecular differences with as-yet unclear functional implications^15, 53^ (called “eccentric” SPNs versus canonical SPNs). We found that the SPN clusters with the strongest enrichment for schizophrenia heritability expressed markers of an eSPN identity, such as *Casz1*, *Htr7*, and *Col11a1*, and were found within both the dorsal and ventral striatum, as well as other striatal and pallidal structures (Fig. 5d,e). Together, these results lend additional support to the potential importance of corticostriatal circuitry in the pathogenesis of schizophrenia, and highlight the value of a brain-wide atlas for nominating disease-relevant cell types.

## Discussion

Here, we combined snRNA-seq and high-resolution spatial transcriptomics with Slide-seq to generate a comprehensive inventory of cell types across each region of the mouse brain. In total, we identified 4,998 clusters of cells, mostly neuronal, with the diversity distributed primarily in subcortical areas, most especially in the midbrain, pons, medulla, and the hypothalamus. We utilized the data to uncover specific neuropeptide signaling interactions, leveraging the specificity of several neuropeptides and/or their receptors. We also characterized activity-related gene expression patterns across all cell types, identifying conserved genes associated with activity, as well as activity-related genes that are more specific to subtypes of neurons. Finally, we nominated specific cell types that are preferentially enriched for the expression of genes associated with human neurological and psychiatric diseases.

A comprehensive inventory of mouse brain cell types should find numerous other immediate uses. One major implication of our analyses is that a substantial fraction of cell types we define are largely unstudied by modern neuroscience methods. To facilitate their interrogation, we deployed an algorithm to identify the minimal set of genes able to specifically define each of our 4,998 clusters. We hope these genes provide a clear path towards the development of genetic tools that can access a wider portion of the astonishing diversity of the nervous system. Beyond achieving more comprehensive access to brain cell types, we anticipate our dataset will drive computational innovations that better neuroanatomically partition the nervous system, and that can integrate other important features of cell type identity such as connectivity, morphology, and physiology. Finally, we expect that our atlas will provide a useful scaffold for interpreting and contextualizing the cell types that are discovered by similar efforts to construct cellular inventories of the human brain^54^.

To facilitate these kinds of applications across neuroscience, we have built a portal to visualize, interact with, and download these data (www.braincelldata.org). Functions have been implemented to plot gene expression and co-expression in CCF-registered space, and within each cell type, and to identify genes and cell types enriched within particular brain regions. We also enable the visualization of spatial localizations of each cell type to specific neuroanatomical structures, and provide a list of minimum marker genes needed to uniquely distinguish them. We hope that facile access to, and interaction with, these rich datasets will provide a firm foundation for functionally characterizing the extraordinarily diverse set of cell types that compose the mammalian brain.

## Acknowledgements

We thank Gordon Fishell, Tushar Kamath, Wade Regehr, and Joshua Welch for helpful discussions. We thank Jackie Goldstein, Daniel King, and the rest of the Hail Batch team for giving computational assistance. This work was supported by NIH/NIMH Brain Grants 1U19MH114821 to E.Z.M. and RF1MH124598 to F.C. and E.Z.M, as well as the Stanley Center for Psychiatric Research.

## Author contributions

E.Z.M. and F.C. conceived the study. J.L. and N.S.S. led the analyses with help from V.G and D.M.C.. J.T.W. developed the snRNA-seq clustering algorithm. M.R. built the data portal, and aligned the adjacent Nissl images to the Slide-seq array data. D.T., C.M., and X.L. performed the CCF integration under supervision from P.M. N.M.N. and C.V. led the snRNA-seq data generation, with help from T.N. K.S.B. led the Slide-seq data generation, with help from E.M. and C.V., under supervision of F.C. J.L., N.S.S., and E.Z.M. wrote the paper, with contributions from all authors.

## Methods

### Animal housing

Animals were group housed with a 12-hour light-dark schedule and allowed to acclimate to their housing environment for two weeks post arrival. All procedures involving animals at MIT were conducted in accordance with the US National Institutes of Health Guide for the Care and Use of Laboratory Animals under protocol number 1115-111-18 and approved by the Massachusetts Institute of Technology Committee on Animal Care. All procedures involving animals at the Broad Institute were conducted in accordance with the US National Institutes of Health Guide for the Care and Use of Laboratory Animals under protocol number 0120-09-16.

### Brain preparation

At 56 days of age, C57BL/6J mice were anesthetized by administration of isoflurane in a gas chamber flowing 3% isoflurane for 1 minute. Anesthesia was confirmed by checking for a negative tail pinch response. Animals were moved to a dissection tray and anesthesia was prolonged via a nose cone flowing 3% isoflurane for the duration of the procedure. Transcardial perfusions were performed with ice cold pH 7.4 HEPES buffer containing 110 mM NaCl, 10 mM HEPES, 25 mM glucose, 75 mM sucrose, 7.5 mM MgCl2, and 2.5 mM KCl to remove blood from brain and other organs sampled. For use in regional tissue dissections, the brain was removed immediately and frozen for 3 minutes in liquid nitrogen vapor and then moved to −80°C for long term storage. For use in generation of the Slide-seq data set via serial sectioning, the brains were removed immediately, blotted free of residual liquid, rinsed twice with OCT to assure good surface adhesion, and then oriented carefully in plastic freezing cassettes filled with OCT. These cassettes were vibrated in a Branson sonic bath for five minutes at room temperature to remove air bubbles and adhere OCT well to the brain surface. The brain’s precise orientation in the X-Y-Z axis was then reset just before freezing over a bath of liquid nitrogen vapor. Frozen blocks were stored at −80°C.

### Construction of brain-wide snRNA-seq dataset

#### Regional dissections

Frozen mouse brains were securely mounted by the cerebellum or by the olfactory/frontal cortex region onto cryostat chucks with OCT embedding compound such that the entire anterior or posterior half (depending on dissection targets) was left exposed and thermally unperturbed. Dissection of anterior-posterior (A-P) spans of the desired anatomical volumes were performed by hand in the cryostat using an ophthalmic microscalpel (Feather safety Razor #P-715) precooled to −20°C and donning 4x surgical loupes. In order to microanatomically assess dissection accuracy, 10 μm coronal sections were taken at relevant A-P dissection junctions and imaged following Nissl staining. Each excised tissue dissectate was placed into a pre-cooled 0.25 ml PCR tube using pre-cooled forceps and stored at −80°C. Nuclei were extracted from these frozen tissue dissectates within 2 days using gentle, detergent-based dissociation as described below.

#### Generation of nuclei suspension and construction of snRNA-seq libraries

Nuclei were isolated from regionally dissected mouse brain samples as previously described^7, 55^. All steps were performed on ice or cold blocks and all tubes, tips, and plates were precooled for >20 min before starting isolation. Dissected frozen tissue in the cryostat was placed in a single well of a 12-well plate and 2 ml of extraction buffer (Ext buffer) was added to each well. Mechanical dissociation was performed by trituration using a P1000 pipette, pipetting 1 ml of solution slowly up and down with a 1-ml Rainin tip (no. 30389212), without creation of froth or bubbles, a total of 20 times. The tissue was allowed to rest in the buffer for 2 min and trituration was repeated. In total, four or five rounds of trituration and rest were performed. The entire volume of the well was then passed twice through a 26-gauge needle into the same well. Approximately 2 ml of tissue solution was transferred into a 50-ml Falcon tube and filled with wash buffer (WB) for a total of 30 ml of tissue solution, which was then split across two 50-ml Falcon tubes (!15 ml of solution in each tube). The tubes were then spun in a swinging-bucket centrifuge for 10 min at 600*g* and 4 °C. Following spinning, the majority of supernatant was discarded (!500 μl remaining with the pellet). Tissue solutions from two Falcon tubes were then pooled into a single tube of !1,000 μl of concentrated nuclear tissue solution. DAPI was then added to the solution at the manufacturer’s (Thermo Fisher Scientific, no. 62248) recommended concentration (1:1,000). Following sorting, nuclei concentration was counted using a hemocytometer before loading into a 10X Genomics 3’ V3 Chip.

#### snRNA-seq library preparation and sequencing

The 10X Genomics (v.3) kit was used for all single-nuclei experiments according to the manufacturer’s protocol recommendations. Library preparation was performed according to the manufacturer’s recommendation. Libraries were pooled and sequenced on NovaSeq S2.

### Construction of brain-wide Slide-seq dataset

#### Generation of larger surface area Slide-seq arrays

Slide-seq arrays were generated as previously described^20^ with slight modifications. Larger diameter gaskets were used to generate 5.5 mm x 5.5 mm, 6.0 mm x 6.2 mm and 6.5 mm x 7.5 mm bead arrays. These sizes were chosen to facilitate different anterior to posterior coronal section sizes. To facilitate image processing, we utilized 2 x 2 digital binning on the collected data, resulting in 1.3 μm per pixel.

#### Serial sectioning procedure

An OCT embedded P56 wild type female mouse brain was thermally equilibrated in the cryostat at −20°C for 30 minutes and then mounted precisely such that an accurate anatomical alignment was maintained. Just anterior to the end of the olfactory bulb region, a 10 μm-thick coronal slice was set as a starting slide. This starting slide was marked, and the following adjacent 10 μm section was used for Slide-seq library preparation. For each tissue slice used for Slide-seq, a 10 μm pre and 10 μm post slide was collected for histology. These histology slides were Nissl stained according to our previously released protocol^56^. After each 10 μm post slice, an 80 μm gap was trimmed before the next set of serial sections was collected; making each Slide-seq slide interval 100 μm apart. A total of 114 sets of three consecutive slides were collected. All pre and post slides for histology registration were stored at −80°C until the slides were Nissl stained. Optimizations were performed in order to be able to hold the Slide-seq tissue slices frozen onto their respective pucks at −80°C during the two days required to complete serial sectioning.

#### Library generation and sequencing

Following the serial sectioning procedure, to process multiple samples at the same time, 10 μm thick tissue slice sections were melted onto Slide-seq arrays and stored at −80°C for two days. On the third day, the frozen tissue sections on the puck were thawed and transferred to a 1.5 mL tube containing hybridization buffer (6X sodium chloride sodium citrate (SSC) with 2 U/μL Lucigen NxGen RNAse inhibitor) for 30 minutes at room temperature. To generate libraries, Slide-seqV2 protocol was adapted from the previously published Slide-seqV2 protocol^20, 57^, in which the volume of reagents was scaled to accommodate the larger surface array of the arrays. Libraries were sequenced using the standard Illumina protocol. The samples were sequenced on either NovaSeq 6000 S2 or S4 flowcells, at a depth of 1.1 – 1.5 billion reads per array, adjusting for the array size. Samples were pooled at a concentration of 4 nM and followed the read structure previously described^20^.

#### Imaging of Nissl sections

We acquired Nissl images on an Olympus VS120 microscope using a 20x 0.75NA objective. Images were captured with a Pike 505C VC50 camera, under auto-exposure mode with a halogen lamp at 92% power. The pixel size in all images was 0.3428 μm, in both the height and width directions. We acquired a total of 114 Nissl images, each from an adjacent section of the brain to a corresponding section that was processed using the Slide-seq pipeline. Of the 114 sections, we removed 10–from the posterior medulla and upper spinal cord–that were outside of the area of the CCF reference brain. Of the remaining 104 images, we removed an additional 3 sections because of unsatisfactory quality of the corresponding Slide-seq puck data. The remaining 101 images comprise the final dataset that we use for all our analyses.

### Registration of Slide-seq data to common coordinate framework

#### Alignment of Slide-seq arrays to adjacent Nissl sections

As a preprocessing step for the alignment of Slide-seq arrays to Nissl images, for each puck, we generated a grayscale intensity image from the Slide-seq data by summing the UMI counts (across all genes) at each bead location on the puck and normalizing by the maximum UMI count value across the entire puck. We then performed the alignment of these images to the adjacent Nissl images in two steps. First, we transformed each Nissl image to an intermediate coordinate space using a manual, rigid transformation. The purpose of this first transformation is to bring all the Nissl images to an approximately equivalent, upright orientation, which made the second step of alignment easier. In the second step, we manually identified corresponding fiducial markers in the Nissl images and Slide-seq intensity images using Slicer3D tool^58^ along with the IGT fiducial registration extension^59^. We then computed the bead positions for all beads via thin plate spline interpolation, where the spline parameters were determined using the fiducial markers.

#### Alignment of Nissl sections to the CCF

Our series of Nissl sections, downsampled to 50 μm resolution by local averaging, were aligned to the 50 μm CCF by jointly estimating 3 transformations. First, a 3D diffeomorphism modeled any shape differences between our sample and the atlas brain. This transformation is modeled in the Large Deformation Diffeomorphic Metric Mapping (LDDMM) framework^60^. Second, a 3D affine transformation (12 degrees of freedom) modeled any pose or scale differences between our sample and the deformed atlas. Third, a 2D rigid transformation (3 degrees of freedom per slice) on each slice modeled positioning of samples onto microscopy slides.

Dissimilarity between the transformed atlas and our imaging data was quantified using an objective function we developed previously^61, 62^, equal to the weighted sum of square error between the transformed atlas and our dataset, after transforming the contrast of the atlas to match the color of our Nissl data at each slice. To transform contrasts, a 3rd order polynomial was estimated on each slice of the transformed atlas to best match the red, green, and blue channels of our Nissl dataset (12 degrees of freedom per slice). During this process, outlier pixels (artifacts or missing tissue) are estimated using an expectation maximization algorithm, and the posterior probabilities that pixels are not outliers are used as weights in our weighted sum of square error.

This dissimilarity function, subject to LDDMM regularization, is minimized jointly over all parameters using a gradient-based approach, with estimation of parameters for linear transforms accelerated using Reimannian gradient descent as recently described^63^. Gradients were estimated automatically using pytorch, and source code for our standard registration pipelines are available online at https://github.com/twardlab/emlddmm. The transformations above were used to map annotations from the CCF onto each slice. The boundaries of each anatomical region were rendered as black curves, and overlaid on the imaging data for quality control (QC). We visually inspected the alignment accuracy on each slice and identified 15 outliers, where our rigid motion model was insufficient due to large distortions of tissue slices. For these slices we included an additional 2D diffeomorphism to model distortions that are independent from slice to slice and cannot be represented as a 3D shape change, as in our previous work^64^. Extended Data Fig. 2a shows accuracy before and after applying the additional 2D diffeomorphism.

#### Analysis of CCF accuracy

We analyzed three genes with highly stereotyped and regional expression: *Dsp*, *Ccn2*, and *Tmem212* which correspond to the following CCF regions:

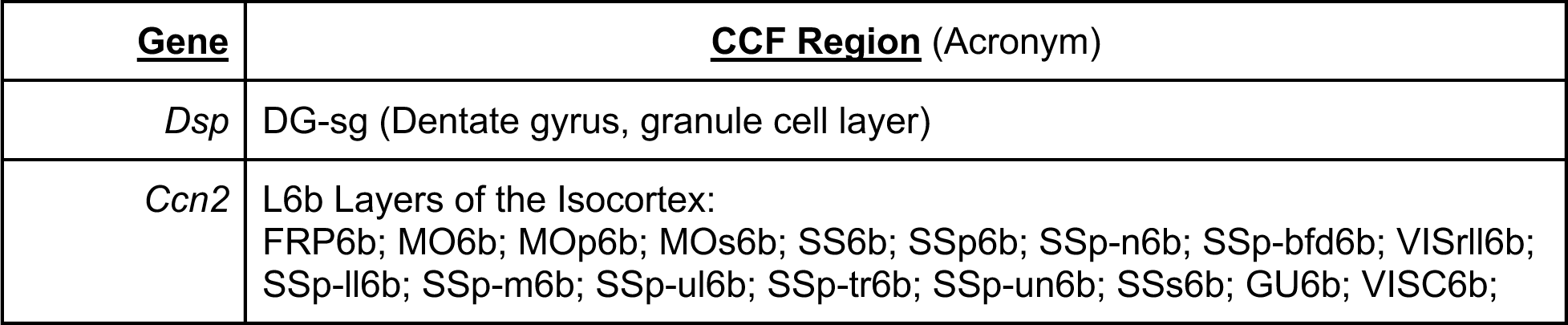
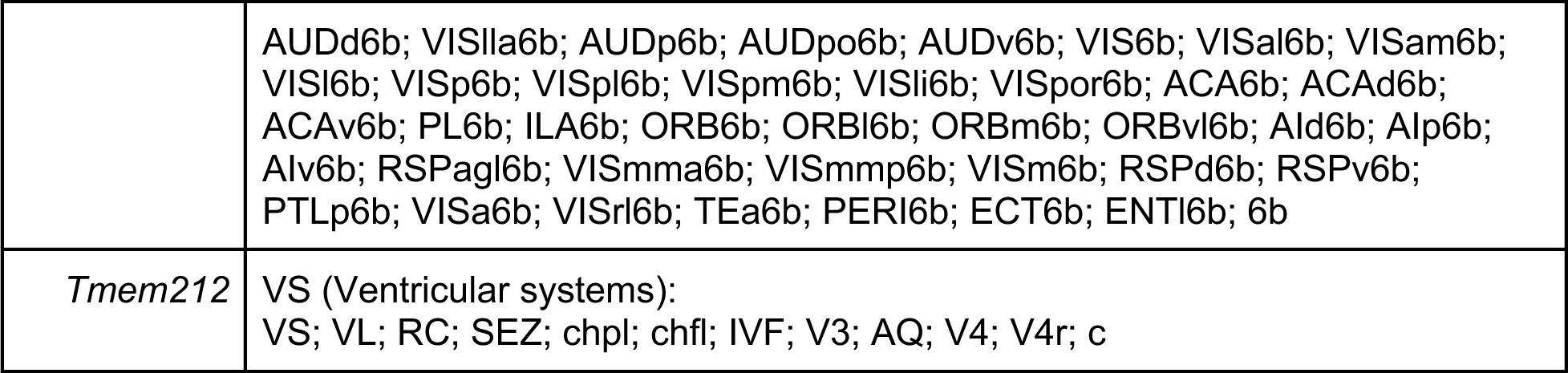

For each bead with non-zero expression of the specified genes, we calculated the distance to the corresponding CCF regions. For preliminary quality control, we used the dbscan package^65^ with eps=3 to filter the points, and used the full width at half maximum metric to summarize the distances (Extended Data Fig. 2c).

### Clustering of snRNA-seq data

#### Overview

Clustering was performed hierarchically, starting from the full dataset of ∼6 million single nuclei. Each round of clustering consisted of: 1) gene selection based on a binomial model; 2) square-root transformation of the counts; 3) construction of the k nearest neighbor and shared neighbor graphs; and 4) Leiden clustering over a range of resolution parameters to find the lowest resolution that yielded multiple clusters. The resulting clusters were then each iteratively re-clustered, and the process was repeated until either: a) no Leiden resolution resulted in a valid clustering or b) the resulting clusters did not have at least three differentially expressed genes distinguishing them. A key goal of this clustering strategy was to re-calculate gene selection for every clustering, as the relevant variable genes depend on the overall context of the cells being clustered. This resulted in a distributed design in which the data was stored on disk in a compressed representation that could be efficiently accessed using parallel processes. This allowed us to perform clustering thousands of times without creating redundant copies of the data.

#### Variable gene selection

To identify variable genes, we used a binomial model of homogenous expression, and looked for deviations from that expectation, similar to a recently described approach^66^. Specifically, for each gene we computed the relative bulk expression by summing the counts across cells and dividing by the total UMIs of the population. This is the proportion of all counts that are assigned to that gene. We use this value as *p* in a binomial model for observing the gene in a cell with *n* counts (equivalently, *np* is equivalent to *λ* in a Poisson model). The expected proportion with non-zero counts is thus:

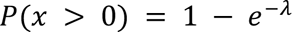

We compared this expected value to the observed percentage of non-zero counts, and selected all genes that are observed at least 5% less than expected in a given population.

#### Construction of shared nearest neighbor graphs

After selecting variable genes, we constructed a shared nearest neighbor (SNN) graph^67, 68^. First, we transformed the counts with the square root function, and then computed the k-nearest neighbor (kNN) graph using cosine distance and *k* = 50 (not including self-edges). From the kNN graph we compute the shared neighbor graph, where the weight between a pair of cells is the Jaccard similarity over their neighbors:

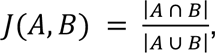

where A and B represent the sets of neighbors for two cells in the kNN graph.

#### Leiden clustering

Once we computed the SNN graph, we used the Leiden algorithm to identify cell clusters, using the Constant Potts Model for modularity^69^. This method is sensitive to a resolution parameter, which can be interpreted as a density threshold that separates inter-cluster and intra-cluster connections. To find a relevant resolution parameter automatically, we implemented a sweep strategy. We started with a very low resolution value, which results in all cells in one cluster. We gradually increased the resolution until there were at least two clusters and the size ratio between the largest and second-largest cluster was at most 20, meaning that at least 5% of the cells are not in the largest cluster. Any cluster of fewer than 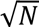 cells was discarded, where *N* was the number being clustered in that round. This discarded set constituted roughly 1.6% of the total cells (100,280 out of 5.9M).

#### Clustering termination and marker gene search

The clustering strategy described above was applied recursively on the leaves of the tree until one of the following conditions was met:

- If the shared neighbor graph was not a single connected component, there is no resolution low enough to form a single cluster, and so the resolution sweep was not possible. This would typically occur if there were very few variable genes, which is indicative of a homogenous cell population.
- If the resolution sweep concluded at the highest resolution without ever finding multiple clusters, this is also indicative of a homogenous population and clustering was considered completed.
- Finally, we truncated the tree when the resulting clusters did not have differentially expressed markers that defined them.

To test for differential markers, we considered each leaf versus its sibling leaves. We used a Mann-Whitney U test to assess whether any genes are differentially expressed. As an additional filter, we required that a gene be observed in less than 10% of the lower population, and observed at a rate at least 20% higher in the higher population, to ensure that there is a discrete difference in expression between the two populations. We required every cluster to have at least three marker genes distinguishing it from its neighbors, as well as three marker genes in the other direction. If a cluster failed that test, all leaves were merged and the parent was considered the terminal cluster.

The only exception to the above was if the next level of clustering resulted in a set of differential clusters that passed this test–these were situations where the first round of clustering split the cells on a continuous difference in expression but the next round resolved the discrete clusters. We retained these clusters for further sub-clustering as they may contain additional structure.

#### Visualization of clusters

For high-dimensional visualization, as in Fig. 1a, we first subsetted each of the clusters to a maximum of 2,000 nuclei. We then performed t-SNE with the Seurat package using 250 principal components over a shared nearest-neighbor graph with a 20 neighbor local neighborhood.

#### Visualization of Cluster Gene Expression

For the heatmap visualization in Fig. 1c, we subsetted the 1,937 mapped cell types to the 1,260 neuronal cell types with at least 5 confidently mapped beads in at least one puck. We normalized the data with Seurat’s LogNormalize normalization (scale.factor=1e4) and averaged each cell type’s 5 nearest neighbors’ expressions. The main region assignment was determined by combining the 10 nearest neighbors’ imputed main region assignment. The matrix was plotted using the ComplexHeatmap package in R^70^.

#### Quality control of clusters

A strict, multi-step quality assessment framework was employed to retain only high quality cell profiles in our analyses. First, we removed nuclei with less than 500 UMIs and greater than 1% mitochondrial UMIs. Doublet clusters were further flagged and excluded based on co-expression of marker genes of distinct cell classes (e.g. *Mbp* and *Slc17a7*).

Next, we constructed a cell “quality-network” to systematically identify and remove remaining low quality cells and artifacts from the dataset. By simultaneously considering multiple quality metrics, our network-based approach has increased power to identify low quality cells while circumventing the issues related to setting hard thresholds on multiple quality metrics. To construct the quality-network we considered the following cell-level metrics: 1) percent expression of genes involved in oxidative phosphorylation (OXPHOS), 2) percent expression of mitochondrial (MT) genes, 3) percent expression of genes encoding ribosomal proteins, 4) percent expression of immediate early gene (IEG) expression, 5) percent expression explained by 50 highest expressing genes, 6) percent expression of long non-coding RNAs (lncRNA), 7) number of unique genes (nGene) (log2-transformed), and 8) number of unique UMIs (nUMI) (log2-transformed). Given their inherently distinct distributions of quality metrics, we separately constructed quality-networks for neurons and glial cells. The quality-network was constructed and clustered using shared-nearest neighbor and Leiden clustering (resolution 0.8) algorithms from Seurat v4.2.0. Our strategy was to remove any cluster from the quality-network with “outlier” distribution of quality metric profiles. A distribution of quality metric was considered as an outlier if its median was above 85% of cells in three features of the quality-network: OXPHOS, MT, and ribosomal protein expression. We further removed any remaining clusters with <15 cells.

#### Estimation of snRNA-seq sampling depth

We used the R package SCOPIT v1.1.4^71^ to estimate the sequencing saturation of our dataset. Under the prospective sequencing model, SCOPIT calculates the multinomial probability of sequencing enough cells, *n\**, above some success probability, *p\**, in a population containing *k* rare cell types of size *N* cells, from which we want to sample at least *c* cells in each cell type:

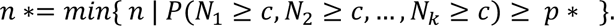

We assume there are *k*=19 rare cell types in our population, each containing *N*=101 cells (frequency of 0.0024%). We need to sequence at least *c*=81 cells from each cell type for sufficient sampling (80% of the rarest cell type). We used SCOPIT to estimate the sampling saturation of our full dataset of 4,388,420 cells.

### Cell type mapping into the Slide-seq dataset with RCTD

We used RCTD to map the single-nuclei clusters onto the Slide-seq spatial beads. For mapping we deployed a modification of the RCTD algorithm^72^ in which we increased the computational efficiency and throughput, modified cell type prefiltering, and adjusted the metric used for the decomposition assignment (see below).

#### Changes to RCTD for parallelizable throughput

We changed the quadratic programming optimizer of RCTD to OSQP^73^ which scales better for the larger matrices resulting from larger sets of cell types to be mapped. We also rewrote the inner loops of the most time-intensive functions (choose_sigma_c and fitPixels) with Rcpp^74^ for efficiency. Additionally, we used Hail Batch^75, 76^ and GNU Parallel^77^ which allowed for large-scale, on-demand parallelization (to thousands of cores) using cloud computing services.

#### Changes to RCTD for cell-type prefiltering

RCTD in doublet mode models how well explicit pairs of cell types match a bead’s expression. For computational efficiency, RCTD prefilters which cell type pairs are considered per bead. However, we found that larger cell type references with many similar cell types led to overly sparse prefiltering which impeded our ability to confidently map fine-grained cell types. In order to balance this sparsity, we added an additional ridge regression term to RCTD’s quadratic optimization, tunable with a ridge strength parameter, which allowed us to control the relative sparsity and potential overfitting of the prefiltering stage. Our modified prefiltering stage used a heuristic to detect a subset of potential cell types for each bead by using RCTD’s full mode with two ridge strength parameters (0.01, 0.001), as well as mapping each cell type individually.

In accordance with the explicit cell type pairs used within RCTD’s doublet mode, we subdivided this filtered list, pulling out the ten cell types deemed most likely to be associated with the given bead. When modeling how well these cell types mapped to a given bead, we exhaustively used one cell type from the top-ten list, and one cell type from the rest of the prefiltered list. For the cerebellum and striatum, the number of cell types considered was sufficiently low that we were able to run the algorithm using all pairs.

#### Changes to RCTD for decomposition assignment

To aid in mapping large references with many similar clusters, we modified how RCTD scores explicit pairs of cell types in doublet mode. Rather than using the result of the single cell type pair which fit best, we identified the cell type pairs which scored similar to the best-scoring pair (with likelihood score within 30). Then, we collated the frequency of each cell type occurring in these well-fitting pairs and divided by the total occurrences of all the cell types, to make a confidence score. Throughout the paper, we use 0.3 (out of a maximum score of 0.5) as the threshold for a ‘confident’ mapping.

#### Creation of per-region cell type references and gene Lists

In order to help reduce the computational load of combinatorially mapping the cell types to each bead, we created a set of tailored references for each region. First, we grouped the libraries into at least one of 8 large-scale regions corresponding to: 1) the basal ganglia, 2) medulla and pons, 3) cerebellum, 4) hippocampal formation, 5) isocortex, 6) midbrain, 7) olfactory bulb, and 8) striatum. For each reference region, the clusters used for mapping had a minimum of 50 cells from the aforementioned per-region libraries and at least 100 cells total.

For each reference region we also generated a tailored gene list. First, for each cluster in each reference region, we ran the same Mann-Whitney U test as in the cluster generation (see above), where the background expression was the other clusters in the reference set. Then we combined all results per gene and chose the 5,000 genes with the smallest p-value across all the individual differential expression tests.

#### Running RCTD on per-region puck subsets

We assigned the CCF regions into at least one of the 8 large-scale regions from above. Then for each Slide-seq puck, we grouped the beads on the puck into at least one of the large-scale regions using our CCF alignment. For each large-scale region on each puck, we ran RCTD using the corresponding tailored reference cell types and tailored gene list. We additionally considered only beads which had at least 150 UMIs across all genes and at least 20 UMIs within the tailored gene list.

### Analyses of cluster heterogeneity across regions

To assess cluster heterogeneity across regions with vastly different areas, we analyzed the minimum number of cell types required to cover 95% of the mapped beads. For each region, we computed the number of confidently mapped beads for each cell type, sorted in descending order by the number of beads. Next, we determined the number of cell types necessary for the running sum of beads to reach 95% of the total mapped beads.

#### Force-directed DeepCCF Region Graph

To generate the force-directed graph of regional cell-type similarity, as in Fig. 2b, we weighted each pair of DeepCCF regions with the weighted Jaccard similarity metric. We then used the R package qgraph v1.9 to generate a force-directed graph.

### Discovery of combinatorial marker genes needed to distinguish snRNA-seq cell types

To find the minimally-sized gene lists which allowed us to distinguish one cell type from the others in the dataset, we framed the question as a set covering problem. In the set cover problem, we find the smallest subfamily of a family of sets that can still cover all the elements in a universe set. For our use case, we examine a given cell type *A* and define a family of sets, one set per gene. The set *S_g_*,corresponding to the gene *g*, contains all the other cell types which are ‘distinguishable’ (defined below) from *A* using the gene *g*. In this way, the minimal set cover of this family of sets gives the smallest subset of genes which distinguishes *A* from all the other cell types in the dataset (i.e. the universe set).

To determine if a gene *g* is ‘distinguishable’ between two cell types *A* and *B*, first we define *Z_A_*_,*g*_ and *Z_B,g_* to be the percent of cells in cluster *A* and cluster *B* with nonzero values of gene *g*, respectively. Then we say *A* is ‘distinguishable’ from *B* using *g*, denoted *D*(*g*, *A*, *B*) = 1, when *Z_A_*_,*g*_ ≥ 0.25 and ((*Z_A_*_,*g*_ – *Z_B,g_*) ≥ 0.5 or *Z_B,g_* ≤ 0.05). Otherwise, we say *A* is “indistinguishable” from B using gene *g*, denoted as *D*(*g*, *A*, *B*) = 0.

Then to solve each cell type’s minimum set cover to optimality (or to prove it is infeasible), we phrase this problem as a mixed integer linear programming (MILP) optimization problem.

We’ll consider one cell type *A* of which we want to find the set cover. Given the *C* other cell types: *c*_1_, *c*_2_,…, *c_c_*; which express *G* genes: *g*_1_, *g*_2_,…, *g_G_*; we create a matrix *M* of dimension (*C* × *G*) where *M_i,j_* = *D*(*g_j_*, *A*, *C_i_*).

We formulate our set cover model as follows:

*Decision variable*. The main variable we want to optimize is the choice of which genes to select for the set cover. So, we define the binary decision variable:

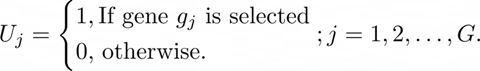

*Constraints*. We then define *X* = *MU*, a *C*-length vector where *X*_*_ gives how many times cell type *c*_*_ was covered by the genes selected in *U*. For *U* to give a proper set cover (i.e. every cell type is covered at least once by the selected genes), we constrain the model such that

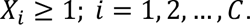

Note that if we want to exclude cell types some cell types *C_a_* and *C_b_* from the optimization (e.g. they are the two nearest neighbors to *A*) we can subset *X* to exclude these two rows. Equivalently, we instead generate a one-hot encoding for the cell types to be excluded with

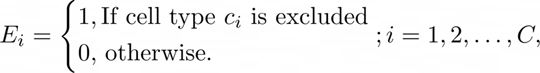

and redefine *X* = (*MU*) + *E*.

*Objective*. Our goal is to minimize the number of genes chosen. Formally

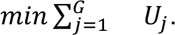

We can define this mixed integer linear model programmatically using the JuMP domain-specific modeling language in Julia^30, 78^. We optimized using the HiGHS open source solver^79^ or the IBM ILOG CPLEX commercial solver v22.1.0.0^80^. We repeated this optimization for every cell type in the dataset. We also group the cell types by main region and repeat this optimization considering only cell types within each regional group.

To enumerate all possible gene lists, for use in obtaining the minimum-sized gene list (below), we used the IBM CPLEX solver with a 5 hour time-limit over two threads and the following parameters:

`mip pool absgap 0`

`mip pool intensity 4`

`mip limits populate 10000000`

### Creation of minimum-sized collated gene list

By default, MILP optimizers will stop after finding one optimal gene list. But, upon examination, we found that most cell types had many distinct equally-optimally-sized gene lists. We again used a set cover approach to leverage this redundancy and determine the smallest possible set of genes that would encompass at least one gene list from each cell type. Using this method we found a ∼25% reduction in the size of the gene list, as opposed to taking the union of first-discovered gene lists returned by the solvers by default.

We first define the *G* expressed genes: *g*_1_, *g*_2_,…, *g_G_*. Using the above exhaustive enumeration using IBM CPLEX, we have a list of equally optimal gene sets for each of the *C* cell types: *c*_1_, *c*_3_,…, *c_c_*. We define *N*_1_, *N*_2_,…, *N_C_* to be how many gene lists each cluster has. In total we have 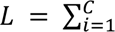 gene lists, enumerated as *l*_1_, *l*_2_,…, *l_L_*. Note that lists (*l*_1_, *l*_2_,…, *l*_N_1__) all come from cell type *c*_1_, lists (*l*_N_1_+1_, *l*_N_1_+2_,…, *l*_N_1_+N_2__) all come from cell type *c*_2_ and so on. For ease of further reference, we will use *i*|*k* to denote the index of the *k*’th gene list for cell type *C_i_* in the whole *L*-length gene list. For example, 1|1 = 1 (the first list of the first cell type is in position 1), *C*|*N*_C_= *L* (the last list of the last cell type is in the final or *L*’th position), and 2|1 = *N*_1_ + 1 (above).

We can encode the individual gene membership for each gene list with a (*L* × *G*) matrix *M* where *M_i,j_* gives whether gene *g_j_* is a member of list *l_i_*.

We define our MILP model such that:

*Decision Variables.* The main binary decision variable we want to optimize is again

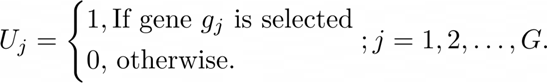

In this case, we need an additional decision variable for each of the *L* gene lists encoding whether it was completely covered by the genes selected in *U*. As we will see, this allows us to require at least one gene list to be completely covered. We define the binary variable

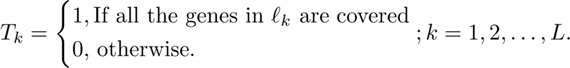

*Constraints.* We define *X* = *MU*, which is a *L*-length vector where *X*_*_ gives how many genes in *l*_*_ are covered by the genes selected in *U*.

For each cell type *C_i_* we’ll create different constraints. Note first that *C_i_* has *N_i_* different gene lists *l_i|1_*, *l_i|2_*,…, *l_i|N_i__* all of the same (optimal) length. We’ll define that length as *S_i_*.

We require at least one of these gene list to be fully covered so constrain

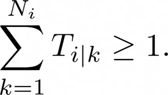

Then we want to enforce the requirements of decision variable *T*. So, for a gene list *l_i|k_*, if all the genes in this set are covered by the chosen genes from *U*, then *T_i|k_* = 1. Using the counterfactual, we see that if *T_i|k_* = 0 then some number of genes in *l_i|k_* are *not* covered by the genes from *U*. In fact, we have an upper bound on this number, because all of the gene lists being examined are of size *S_i_*, so at most *S_i_* genes are missing from being covered by *U* in *T_i|k_*.

Given a list *l_i|x_* where *T_i|x_* = 1, for it actually to be fully covered by the non-zero elements of *U*, we require *X_i|x_* ≥ *S_i_*. For a list *l_i|y_* where *T_i|y_* = 0, since it is already not fully covered by the non-zero elements of *U*, we can ignore its corresponding *X_i|y_*. One way of ignoring this set, while keeping the broader constraint in place for optimization, is by adding the upper bound of missing genes, *S_i_*, since *X_i|y_* + *S_i_* ≥ *S_i_* is true by inspection (*X_i|y_* ≥ 0 by definition). Putting this together we get the constraint

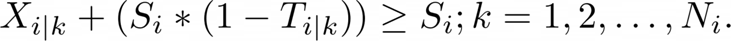

We repeat this constraint for every cluster *C*_*_ and for each cluster grouped by the main region.

#### GO enrichment of genes with hierarchical reduction

The GO enrichment analysis for the minimum-sized collated gene list was performed with enrichr^81, 82^ using the GO Molecular Function 2021 list gene set^83, 84^. This resulting enrichment was hierarchically reduced for visualization with rvvgo^85^.

### Neurotransmitter and neuropeptide assignment to cell types

#### Neurotransmitter assignment

Each cell type was assigned to a neurotransmitter identity based upon the percent of its cells with non-zero counts of genes essential for the function of that neurotransmitter. Specifically, we used a non-zero threshold *nz* = 0.35:

- VGLUT1: *Slc17a7* ≥ *nz*
- VGLUT2: *Slc17a6* ≥ *nz*
- VGLUT3: *Slc17a8* ≥ *nz*
- GABA: (*Gad1* | *Gad2* ≥ *nz*) & (*Slc32a1* ≥ *nz*)
- GLY: (*Gad1* | *Gad2* ≥ *nz*) & (*Slc6a5* | *Slc6a9* ≥ *nz*)
- CHOL: *Slc18a3* & *Chat* ≥ *nz*
- DOP: *Slc6a3* ≥ *nz*
- NOR: *Pnmt* | *Dbh* ≥ *nz*
- SER: *Slc6a4* | *Tph2* ≥ *nz*

For the 166 neuronal cell types that didn’t meet the above *nz* conditions, we carefully examined their top expressing transporters and assigned neurotransmitters accordingly.

#### Neuropeptide assignment

Each cell type was assigned to a neuropeptide (NP) ligand identity if: a) the percent of its cells with non-zero expression of the NP was ≥ 0.3; and b) the average expression of the NP was ≥ 0.5 counts per cell. We observed that the expression of four NPs showed greater contamination across other cell types: OXT, AVP, PMCH, and AGRP. Therefore, for these NPs, we required the percent of cells with non-zero expression to be ≥ 0.8 and average expression to be ≥ 5 counts per cell.

Each cell type was assigned to a neuropeptide receptor (NPR) identity if: a) the percent of its cells with non-zero expression of at least one NPR was ≥ 0.2; b) the average expression of at least one NPR was ≥ 0.5 counts per cell.

### Quantification of region-specific excitatory-inhibitory ratios

We first created inhibitory and excitatory cell type groups based on their neurotransmitter expression, as above. We classified cell types expressing GABA or GLY neurotransmitters as inhibitory and those expressing VGLUT neurotransmitters as excitatory. In the case where a cell type was assigned to both an inhibitory and excitatory identity, it was classified as inhibitory. For each region on the Slide-seq array, we labeled its beads as excitatory or inhibitory by whether they confidently mapped into members of the corresponding cell type groups, with additional filtering to ensure these mappings were one of top 2 ranked cell types per bead. Then, defining #*I* and #*E* gives the number of inhibitory and excitatory mapped beads, respectively, we defined the excitatory-to-inhibitory balance as 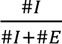.

To quantify the uncertainty, we calculated the exact 95% confidence interval for the corresponding binomial distribution using the exact method of binconf function in the Hmisc R package^86^.

### Analyses of activity-dependent gene expression

#### Pseudocell generation

We aggregated our snRNA-seq expression data into pseudocells: aggregations of cells with similar gene expression profiles. Working at the pseudocell resolution (rather than with individual cells) eliminates the technical variation issues of single cell transcriptomic data, such as low capture rate from dropouts and pseudoreplication through averaging expression of similar cells^87, 88^, while avoiding issues of pseudobulk approaches, such as low statistical power and high variation in sample sizes^89^.

To generate our pseudocells, we first performed dimensionality reduction at the single-cell level. Single cells were divided into 27 groups, consisting of glial cell classes and neuronal populations further divided by neurotransmitter usage. Within each cell group, we selected genes that were highly variable in a specific number of mouse donors, such that a maximum of 5,000 genes would be used for subsequent scaling by batch. We then ran PCA on the scaled expression data (50 PCs for glia and 250 PCs for neurons). Next, we constructed pseudocells by grouping single cells within each cell type. Within a cell type of size *n*, cells were assigned to pseudocells of size *s*, such that the pseudocell size correlated with cell type size:

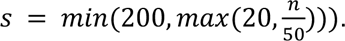

Pseudocell centers were identified by applying k-means clustering on the top PCs (50 for glia and 250 for neurons). To ensure the stability of results across different cell type sizes – ranging from rare neuronal clusters of 15 cells to a glial cluster of half a million cells – we weighted PCs by their variance explained. Random walk approaches have been found to be more robust in identifying cells of similar gene expression profiles, in contrast to spherical, distance-based methods^90, 91^. Therefore, we used the random walk method on cell-cell distances in the PCA space to assign cells to pseudocell centers (i.e. k-means centroids)^91^. To generate our pseudocell counts matrix, we aggregated the raw UMI counts of cells assigned to each pseudocell. This resulted in representation of each cell type by one or more pseudocells, ranging from 1 to 2,490 pseudocells.

The pseudocell-level expression of protein-coding genes was normalized by log2-transformed count per million (CPM), followed by quantile normalization. We further normalized the expression of each gene to have a mean expression of zero and standard deviation of one. Normalized pseudocell counts were used for downstream analysis.

#### Candidate ARG list

To generate our candidate activity-regulated genes (ARGs) list, we divided our neuronal pseudocells into 28 cell groups, such that each main region was assigned to an excitatory and inhibitory population, and all other cell types (cholinergic, dopaminergic, noradrenergic, and serotonergic) were individually grouped across all main regions. To construct gene co-expression networks within each cell group, we computed pairwise gene correlation coefficients (Pearson) across scaled pseudocells using the R package psych v2.2.5. For a gene *g* to be considered an ARG candidate, its correlation *r* with *Fos* must: 1) *r* ≥ 0.3; 2) *r* in the ≥ 99.5% quantile distribution of *g*’s correlations with all genes; and 3) *r* is statistically significant after multiple hypothesis testing (Holm adjusted p-value < 0.05). To construct the final full ARG candidate list, we took the union of selected genes across all cell groups.

#### Activity-regulated gene network

To visualize activity-regulated relationships between our candidate ARGs and regions of the brain, we constructed a force-directed graph of a weighted bipartite network. We used the R package igraph v1.2.7 to build the network from an incidence matrix of candidate ARGs and excitatory/inhibitory cell types localized to different regions. An entry *e* in the matrix corresponds to a gene’s correlation *r* with *Fos* in a brain region, scaled up by 1 such that all entries ≥ 1.The nodes of the network comprised two disjoint sets: candidate ARGs and neuronal brain regions, such that there would never be an edge between a pair of genes or a pair of regions. Edges were weighted based on the correlation entry *e* between a gene and region node. To emphasize the most central nodes in the network, we pruned edges with *e* < 1.3 and after pruning, we removed nodes with degree < 2. Gene node sizes and labels were also adjusted based on the scaled degree of the node to highlight the most central genes. Gene nodes were labeled only if they shared edges of weight *e* ≥ 1.3 with at least six regions. R packages GGally v2.1.2 and ggplot2 v3.3.6 were used to plot the network using the Kamada-Kawai force-directed layout algorithm. We selected core IEGs from the network based on node centrality (degree > 18).

#### Classifying ARG clusters

To further characterize our candidate ARGs, we performed ward.D2 hierarchical clustering based on their *Fos* correlations across brain regions. We cut the dendrogram at a height that divided our ARGs into 7 clusters. To assess the overlap between our ARG clusters and the ARGs reported in Tyssowski et al.^40^, we computed a Fisher’s exact test between two given gene sets using the R package GeneOverlap v1.30.0^92^. P-values were Bonferroni-corrected for multiple hypothesis testing in each gene cluster.

#### Transcription factor enrichment

To identify the transcription factors (TFs) selectively enriched for telencephalic excitatory or inhibitory populations, we performed GSEA on a ranked gene list against a curated TF gene set. First, we ranked genes from our telencephalic excitatory cells by their average correlation with *Fos*, compared to a background ranking of average telencephalic inhibitory *Fos* correlations. Genes from our telencephalic inhibitory cells were ranked in reverse order. We built a gene set of TFs by combining enrichR^82^ databases from ARCHS4, ENCODE, and TRRUST. We subsetted for human TFs that showed up in at least two databases and had at least 10 unique targets in each of those databases. The final gene set consisted of these human TFs, whose targets comprised the intersection between any two databases. We used the R package fgsea v1.20.0^93^ to run GSEA with a gene set size restriction of 15 to 500, and against a background of all protein-coding genes expressed in our normalized pseudocell data.

### Heritability enrichment with scDRS

To determine which of our snRNA-seq cell types were enriched for specific GWAS traits, we used single-cell disease relevance score (scDRS)^47^ with default settings. scDRS operates at the single-cell level to compute disease association scores, while considering the distribution of control gene scores to identify significantly associated cells.

We used MAGMA^94^ to map SNPs to genes (GRCh37 genome build from 1000 Genomes Project) using an annotation window of 10kb. We used the resulting annotations and GWAS summary statistics to calculate each gene’s MAGMA z-score (association with a given trait).

Human genes were converted to their mouse orthologs using a homology database from Mouse Genome Informatics. The 1,000 disease genes used for scDRS were chosen and weighted based on their top MAGMA z-scores. Many of the traits we tested for enrichment had previously computed MAGMA z-scores from Zhang et al.^47^, so those scores were used instead (after applying MGI gene ortholog conversion).

scDRS was used to calculate the cell-level disease association scores for a given trait; in our case, we treated our aggregated raw pseudocell counts as the input single-cell dataset, validating that the pseudocell results largely recapitulated single-cell-level results for three traits (Extended Data Fig. 6a). To determine trait association at the annotated cell type resolution, we used the z-scores computed from scDRS’s downstream Monte Carlo test. These MC z-scores were converted to theoretical p-values using a one-sided test under a normal distribution.Theoretical p-values were FDR-corrected for multiple hypothesis testing, considering only cell types with at least 4 beads confidently mapped to a single puck and deep CCF region, as well as non-neurogenesis cell types.

## Data and Code Availability

An interactive portal is available at www.braincelldata.org, where gene expression data and spatial localizations of individual cell types can be visualized in a number of ways. Raw single-nucleus RNA-seq and Slide-seq data is available in the NeMO Archive (www.nemoarchive.org). The snRNA-seq clustering algorithm is available in the repository www.github.com/MacoskoLab/brain-atlas. This code was built on top of several existing packages: `numpy`, `scipy`, `dask`, `pynndescent`, `leidenalg`, `igraph`, `zarr`, and their dependencies.

## Supplemental Tables

**Supplemental Table 1.** List of snRNA-seq dissectates and their CCF regional compositions.

**Supplemental Table 2.** List of “DeepCCF” regions used in this study.

**Supplemental Table 3.** Minimum gene lists computed to define each cell type, across a range of parameterizations.

**Supplemental Table 4.** List of neuropeptides used in Figure 3, along with their cognate receptors.

**Supplemental Table 5.** List of GWAS traits analyzed in Figure 5.

**Supplemental Table 6.** Cell-type-level enrichment scores (adjusted p-values) computed by scDRS for each GWAS trait.

---

**Extended Data Figure 1.**
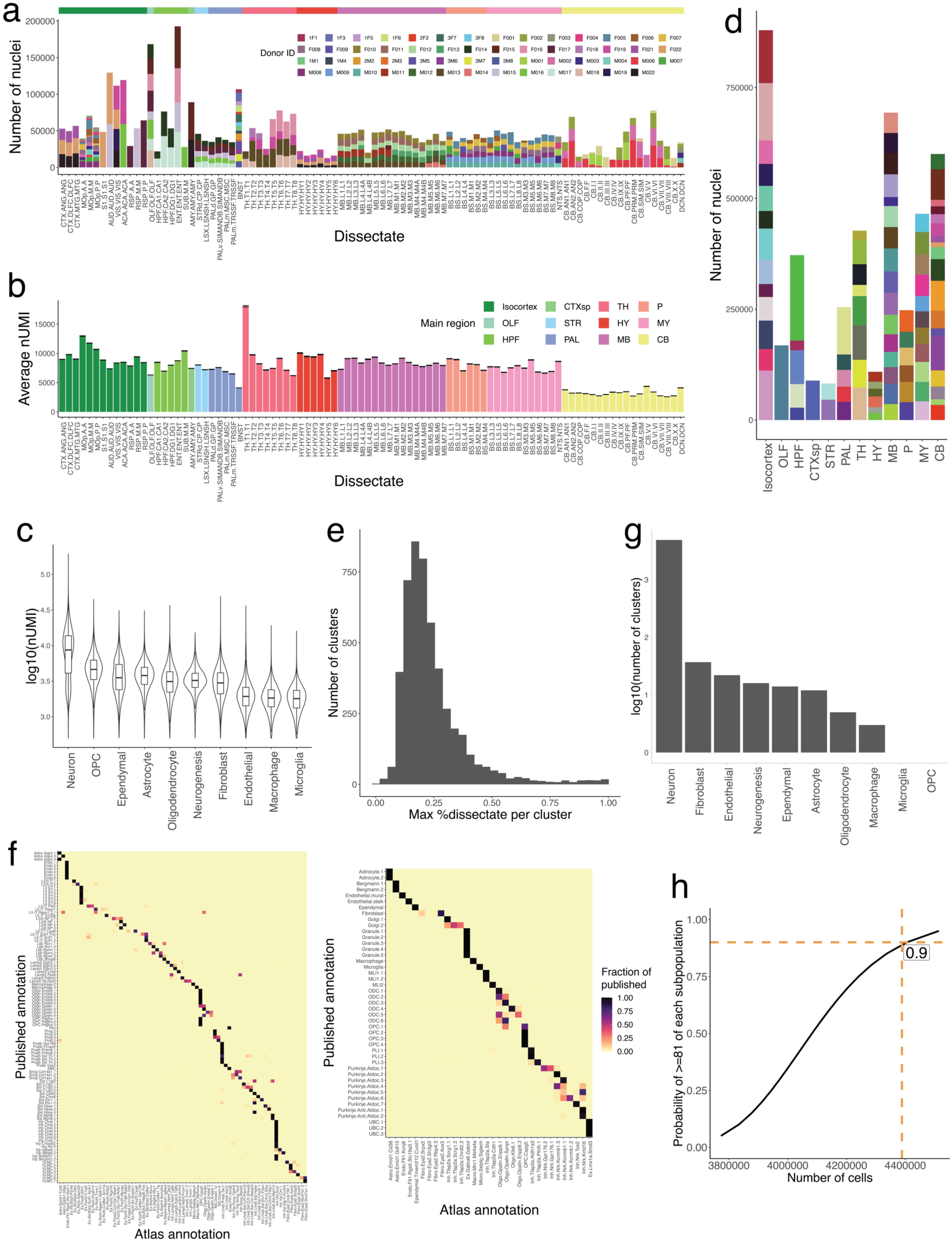
Quality control and summary statistics of the snRNA-seq analysis. **a**, Stacked barplots showing the number of nuclei sampled in each region, for each animal replicate. Female donor IDs contain an “F” in their name, while male donors contain an “M.” Dissectates are colored on the top by their corresponding major brain region. **b,** Bar plots showing the average number of UMIs per nucleus in each dissectate, colored by main brain region. Error bars indicate standard error of each dissectate’s average number of UMIs. **c,** Violin plots showing the log10 distribution of the number of UMIs per nucleus in each major cell class. **d**, Stacked barplots of the number of nuclei sampled in each major mouse brain region, subsetted by individual dissectate. **e**, Histogram of the maximal proportional representation of individual dissectates in each snRNA-seq cluster. **f**, Heatmap representing a confusion matrix between clustering of the snRNA-seq data in the current study (x-axis), and published studies (y-axis), for the mouse motor cortex^4^ (left) and cerebellum^7^ (right). **g**, Histogram of the log10 number of clusters recovered from each major cell class. **h**, Plot indicating the probability of sampling 19 very rare populations (prevalence 0.0024%) as a function of the total number of cells profiled in experiment (probability estimated as in methods). Number of high-quality nuclei profiled here (4,388,420) and corresponding probability are indicated.

**Extended Data Figure 2.**
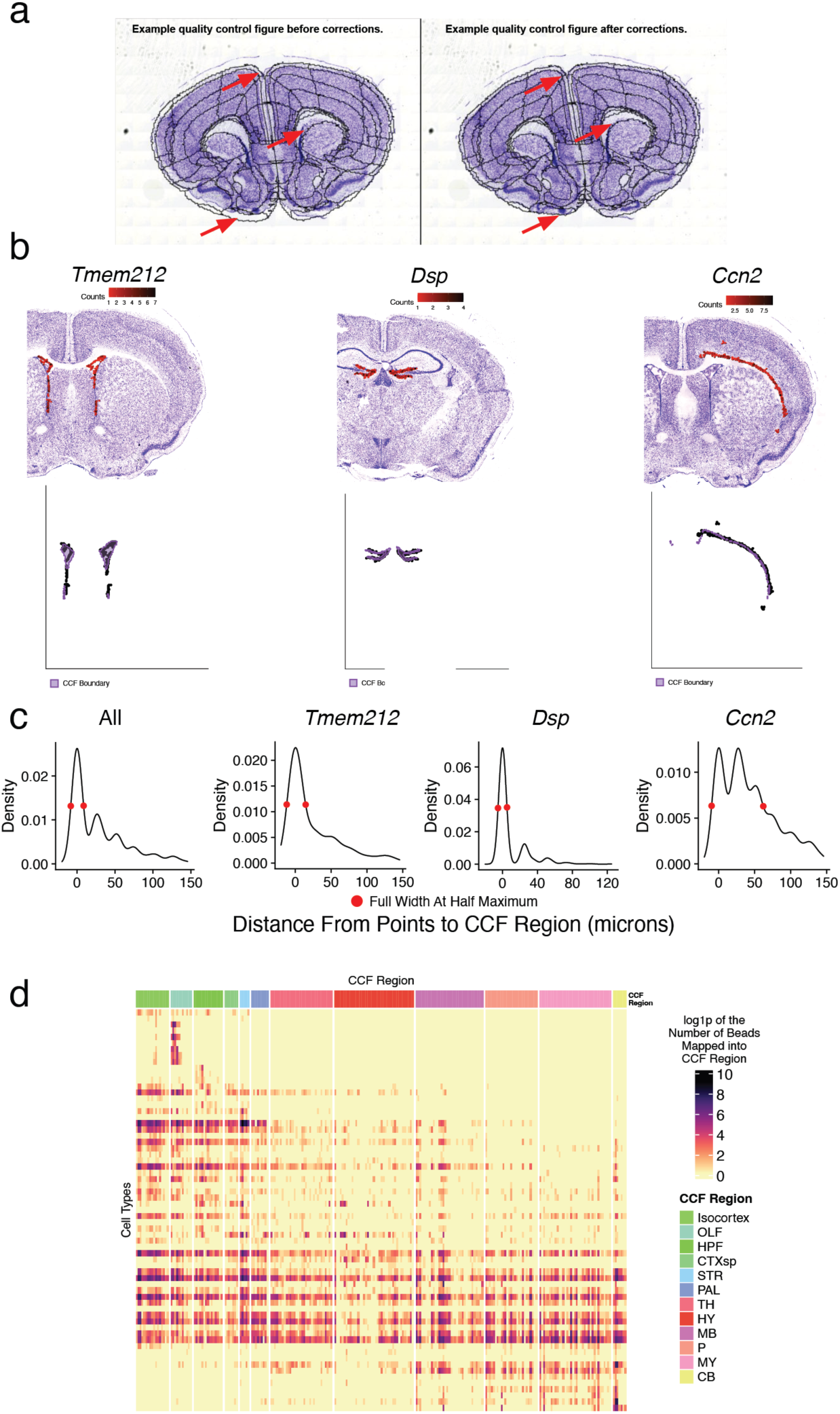
Quality control and summary statistics of CCF integration and cell type mapping. **a,** Example images of adjacent Nissl sections aligned to CCF with 2D rigid transformation (wireframe outline) before (left) and after (right) correcting alignment with a 2D diffeomorphism. Red arrows point to example regions with incorrect alignment, and improvement after application of the correction. **b**, Expression of three highly specific marker genes that label the ventricular lining (*Tmem212*), dentate gyrus granule layer (*Dsp*), and layer 6b of isocortex (*Ccn2*) in Slide-seq (top row). Bottom row shows the positions of individual beads with expression with respect to the boundaries of the expected CCF region (purple). **c,** Density plot of the distance of each bead expressing each of the three marker genes (or all combined) shown in **b** across the corresponding Slide-seq sections. The full width half maximum of the density profile is shown. **d**, Heat map representing the frequency of bead mappings for each glial cell type, across DeepCCF regions.

**Extended Data Figure 3.**
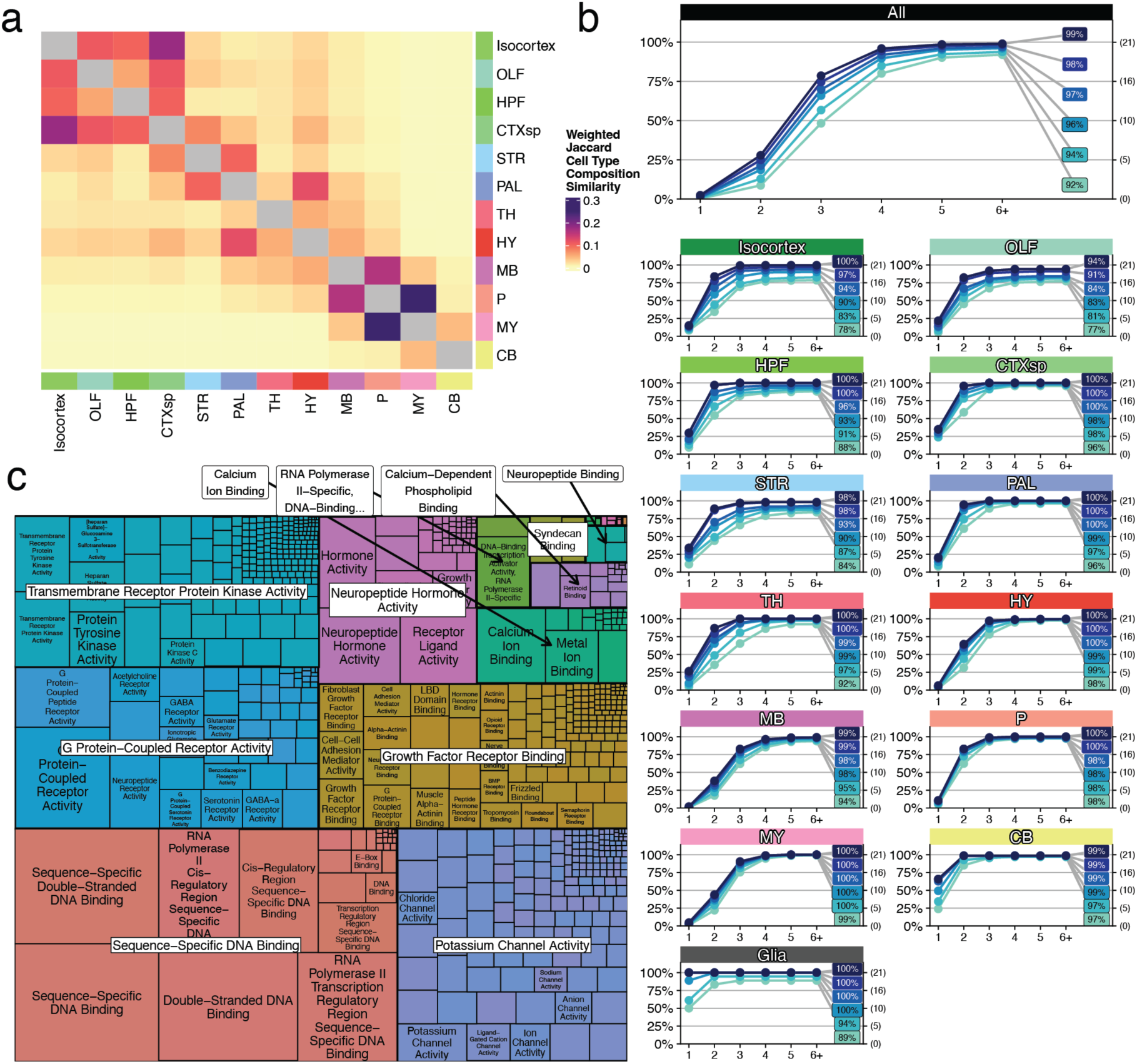
Extended analyses quantifying neuronal cell type diversity across brain areas. **a**, Heatmap representing the weighted Jaccard similarity in cell type composition between each of the 12 main brain areas (Methods). **b**, Cumulative distribution plots of the number of genes needed to label each cell type, within each of the 12 major brain areas. Within each region, each of the sets of individually colored plots denotes an algorithmic run with a different number of nearest neighbors that are tolerated as having the same gene markers (absolute number in parentheses at right). The colored percentages denote the proportion of cell types for which the algorithm was able to find a solution. **c,** Treemap visualization of the GO hierarchy enriched in the minimum-sized collated gene list after hierarchical reduction (Methods).

**Extended Data Figure 4.**
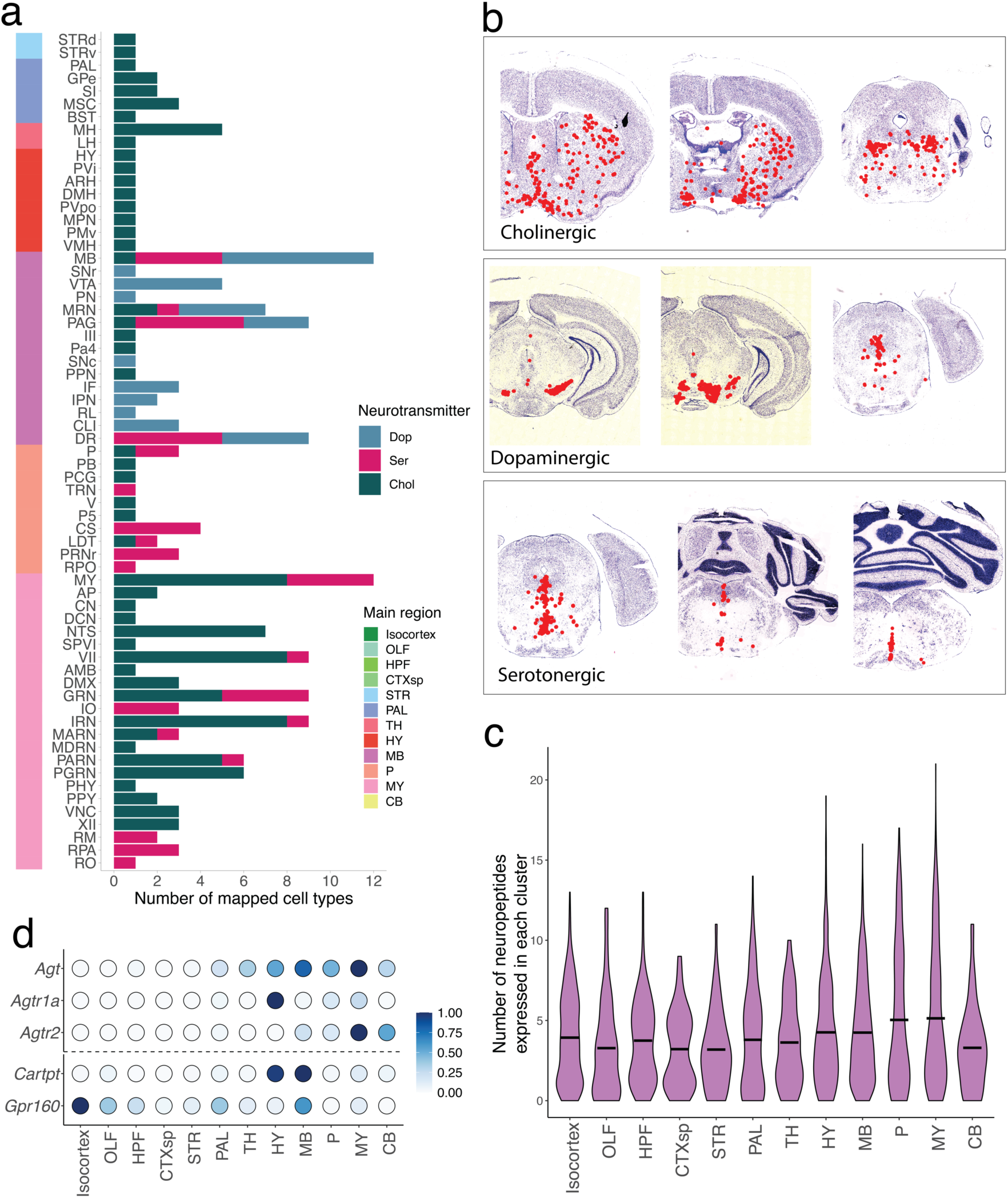
Extended analyses of neurotransmitter and neuropeptide usage across the brain. **a**, Stacked barplots of the number of cell types with confident mappings in each deep CCF region, subsetted by neurotransmitter group. Deep CCF regions are colored on the left by their corresponding major brain region. **b**, Representative sections showing the spatial localizations of all cell types within three neurotransmitter groups. **C**, Violin plots of the number of neuropeptides expressed in each cluster, stratified by main brain region. **d,** Dot plot showing scaled Slide-seq counts per 10,000 of ligand-receptor pairs across main brain regions.

**Extended Data Figure 5.**
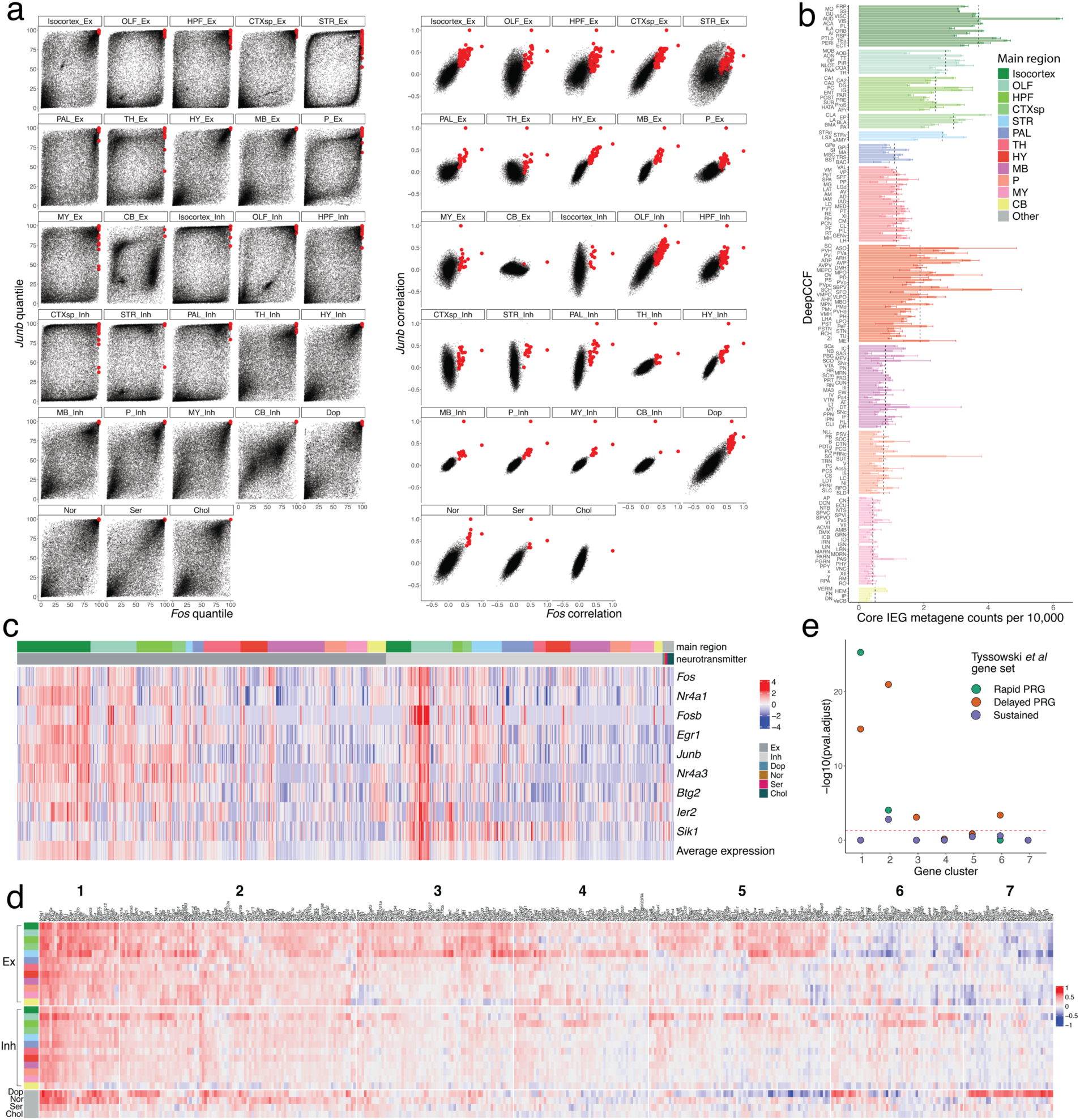
Additional analyses related to activity-related genes. **a,** Comparison of correlations (right) and quantile of correlation (left) between each gene and both *Fos* (x-axis) and *Junb*. Red dots indicate the genes that were selected as candidate ARGs. **b**, Barplots showing the average counts per 10,000 of the core IEG metagenes (Methods) listed in **c**. Error bars indicate standard error of average counts per deep CCF region and dashed black lines indicate average counts per main brain region. **c**, Scaled mean expression of the core IEG metagenes, within each main region, separated by neurotransmitter group. **d**, Extended heat map showing the correlation with *Fos* of all candidate ARGs within the seven clustered groups. **e**, Enrichment analysis (Methods) of each candidate ARG cluster with three established ARG gene sets^40^. Dotted red line indicates an adjusted p-value threshold of 0.05.

**Extended Data Figure 6.**
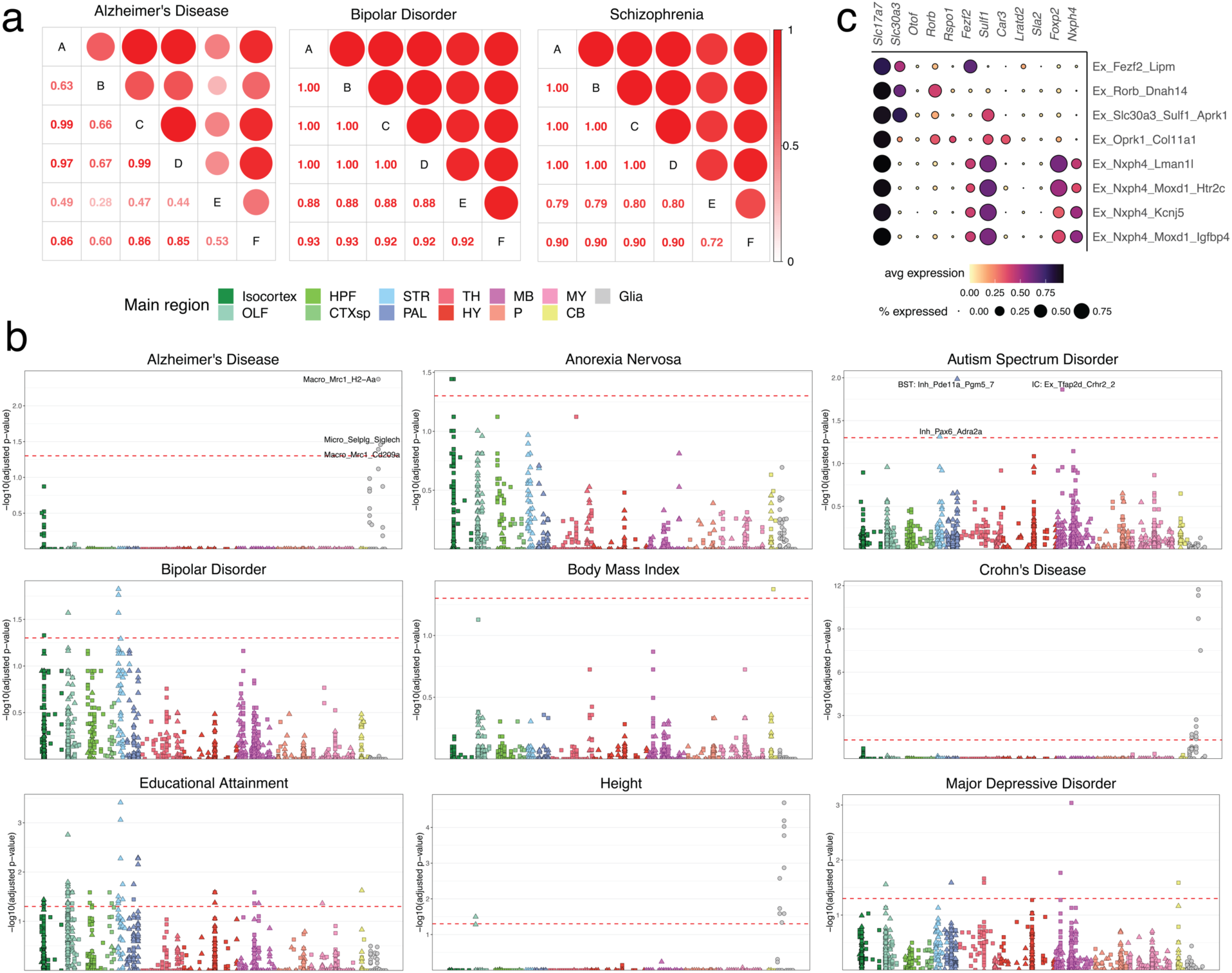
Extended analyses of heritability enrichment in murine brain cell types. **a**, Correlation plot of p-value enrichment scores (FDR-corrected) for each cell type across different scDRS settings: A) default parameters (Methods) B) MAGMA gene z-score > 2.5, C) control gene set size of 2,000, D) control gene set size of 500, E) input expression dataset at the single-cell level (versus pseudocells), F) adjust for cell type proportions. **b,** Adjusted −log10 p-value enrichment scores for each cell type, grouped and colored by main region, for an extended set of GWAS-measured traits. **c**, Dot plot of the expression of key cortical pyramidal cell type markers within the eight isocortical clusters that were significantly enriched for schizophrenia heritability.

## Citations

1. Zeng, H. What is a cell type and how to define it? Cell 185, 2739–2755 (2022).

2. Zeng, H. & Sanes, J. R. Neuronal cell-type classification: challenges, opportunities and the path forward. Nat. Rev. Neurosci. 18, 530–546 (2017).

3. Wang, Q. et al. The Allen Mouse Brain Common Coordinate Framework: A 3D reference atlas. Cell 181, 936–953.e20 (2020).

4. Yao, Z. et al. A transcriptomic and epigenomic cell atlas of the mouse primary motor cortex. Nature 598, 103–110 (2021).

5. Yao, Z. et al. A taxonomy of transcriptomic cell types across the isocortex and hippocampal formation. Cell 184, 3222–3241.e26 (2021).

6. Welch, J. D. et al. Single-Cell Multi-omic Integration Compares and Contrasts Features of Brain Cell Identity. Cell (2019) doi:10.1016/j.cell.2019.05.006.

7. Kozareva, V. et al. A transcriptomic atlas of mouse cerebellar cortex comprehensively defines cell types. Nature 598, 214–219 (2021).

8. Zhang, C. et al. Area Postrema Cell Types that Mediate Nausea-Associated Behaviors. Neuron 109, 461–472.e5 (2021).

9. Campbell, J. N. et al. A molecular census of arcuate hypothalamus and median eminence cell types. Nat. Neurosci. 20, 484–496 (2017).

10. Moffitt, J. R. et al. Molecular, spatial, and functional single-cell profiling of the hypothalamic preoptic region. Science 362, (2018).

11. Okaty, B. W. et al. A single-cell transcriptomic and anatomic atlas of mouse dorsal raphe Pet1 neurons. Elife 9, (2020).

12. Kim, D.-W. et al. Multimodal Analysis of Cell Types in a Hypothalamic Node Controlling Social Behavior. Cell 179, 713–728.e17 (2019).

13. Kebschull, J. M. et al. Cerebellar nuclei evolved by repeatedly duplicating a conserved cell-type set. Science 370, (2020).

14. Zeisel, A. et al. Molecular Architecture of the Mouse Nervous System. Cell 174, 999–1014.e22 (2018).

15. Saunders, A. et al. Molecular Diversity and Specializations among the Cells of the Adult Mouse Brain. Cell 174, 1015–1030.e16 (2018).

16. Ortiz, C. et al. Molecular atlas of the adult mouse brain. Sci Adv 6, eabb3446 (2020).

17. Zhang, M. et al. Spatially resolved cell atlas of the mouse primary motor cortex by MERFISH. Nature 598, 137–143 (2021).

18. Chen, R. et al. Decoding molecular and cellular heterogeneity of mouse nucleus accumbens. Nat. Neurosci. 24, 1757–1771 (2021).

19. Rodriques, S. G. et al. Slide-seq: A scalable technology for measuring genome-wide expression at high spatial resolution. Science 363, 1463–1467 (2019).

20. Stickels, R. R. et al. Highly sensitive spatial transcriptomics at near-cellular resolution with Slide-seqV2. Nat. Biotechnol. 39, 313–319 (2021).

21. Franklin, K. B. J. & Paxinos, G. The Mouse Brain in Stereotaxic Coordinates. (Academic Press, 1997).

22. Dong, H. W. The Allen reference atlas: A digital color brain atlas of the C57Bl/6J male mouse. 57B, 6J (2008).

23. Murray, Zou, Goeva & Macosko. Robust decomposition of cell type mixtures in spatial transcriptomics. Nature.

24. hail: Scalable genomic data analysis. (Github).

25. Kaseniit, K. E. et al. Modular, programmable RNA sensing using ADAR editing in living cells. Nat. Biotechnol. (2022) doi:10.1038/s41587-022-01493-x.

26. Qian, Y. et al. Programmable RNA sensing for cell monitoring and manipulation. Nature 610, 713–721 (2022).

27. Jiang, K. et al. Programmable eukaryotic protein synthesis with RNA sensors by harnessing ADAR. Nat. Biotechnol. (2022) doi:10.1038/s41587-022-01534-5.

28. Karp, R. M. Reducibility among Combinatorial Problems. Complexity of Computer Computations 85–103 Preprint at https://doi.org/10.1007/978-1-4684-2001-2_9 (1972).

29. Chandru, V. & Rammohan Rao, M. Integer Programming. Electronic Journal Preprint at https://doi.org/10.2139/ssrn.2170269.

30. Dunning, I., Huchette, J. & Lubin, M. JuMP: A Modeling Language for Mathematical Optimization. SIAM Review vol. 59 295–320 Preprint at https://doi.org/10.1137/15m1020575 (2017).

31. Characterized cre lines. The Jackson Laboratory https://www.jax.org/research-and-faculty/resources/cre-repository/characterized-cre-lines-jax-cre-resource.

32. van den Pol, A. N. Neuropeptide transmission in brain circuits. Neuron 76, 98–115 (2012).

33. Yosten, G. L. et al. GPR160 de-orphanization reveals critical roles in neuropathic pain in rodents. J. Clin. Invest. 130, 2587–2592 (2020).

34. Kamath, T. et al. Single-cell genomic profiling of human dopamine neurons identifies a population that selectively degenerates in Parkinson’s disease. Nat. Neurosci. 25, 588–595 (2022).

35. Kim, G.-H. J. et al. A zebrafish screen reveals Renin-angiotensin system inhibitors as neuroprotective via mitochondrial restoration in dopamine neurons. Elife 10, (2021).

36. Jo, Y., Kim, S., Ye, B. S., Lee, E. & Yu, Y. M. Protective Effect of Renin-Angiotensin System Inhibitors on Parkinson’s Disease: A Nationwide Cohort Study. Front. Pharmacol. 13, 837890 (2022).

37. Yap, E.-L. & Greenberg, M. E. Activity-Regulated Transcription: Bridging the Gap between Neural Activity and Behavior. Neuron 100, 330–348 (2018).

38. Marsh, S. E. et al. Dissection of artifactual and confounding glial signatures by single-cell sequencing of mouse and human brain. Nat. Neurosci. 25, 306–316 (2022).

39. Hrvatin, S. et al. Single-cell analysis of experience-dependent transcriptomic states in the mouse visual cortex. Nat. Neurosci. 21, 120–129 (2018).

40. Tyssowski, K. M. et al. Different Neuronal Activity Patterns Induce Different Gene Expression Programs. Neuron 98, 530–546.e11 (2018).

41. Luan, S. et al. Thyrotropin receptor signaling deficiency impairs spatial learning and memory in mice. J. Endocrinol. 246, 41–55 (2020).

42. Farrow, P., et al. Auxiliary subunits of the CKAMP family differentially modulate AMPA receptor properties. eLife vol. 4 Preprint at https://doi.org/10.7554/elife.09693 (2015).

43. Glerup, S. et al. SorCS2 is required for BDNF-dependent plasticity in the hippocampus. Mol. Psychiatry 21, 1740–1751 (2016).

44. Bryois, J. et al. Genetic identification of cell types underlying brain complex traits yields insights into the etiology of Parkinson’s disease. Nat. Genet. 52, 482–493 (2020).

45. Skene, N. G. et al. Genetic identification of brain cell types underlying schizophrenia. Nat. Genet. 50, 825–833 (2018).

46. Finucane, H. K. et al. Heritability enrichment of specifically expressed genes identifies disease-relevant tissues and cell types. Nat. Genet. 50, 621–629 (2018).

47. Zhang, M. et al. Polygenic enrichment distinguishes disease associations of individual cells in single-cell RNA-seq data. Preprint at https://doi.org/10.21203/rs.3.rs-933790/v1.

48. Bryois, J. et al. Genetic Identification of Cell Types Underlying Brain Complex Traits Yields Novel Insights Into the Etiology of Parkinson’s Disease. Preprint at https://doi.org/10.1101/528463.

49. Schwartzentruber, J. et al. Genome-wide meta-analysis, fine-mapping and integrative prioritization implicate new Alzheimer’s disease risk genes. Nature Genetics vol. 53 392– 402 Preprint at https://doi.org/10.1038/s41588-020-00776-w (2021).

50. Lichtenstein, P. et al. Common genetic determinants of schizophrenia and bipolar disorder in Swedish families: a population-based study. Lancet 373, 234–239 (2009).

51. Brainstorm Consortium et al. Analysis of shared heritability in common disorders of the brain. Science 360, (2018).

52. Crittenden, J. R. & Graybiel, A. M. Basal Ganglia disorders associated with imbalances in the striatal striosome and matrix compartments. Front. Neuroanat. 5, 59 (2011).

53. Gokce, O., et al. Cellular Taxonomy of the Mouse Striatum as Revealed by Single-Cell RNA-Seq. Cell Reports vol. 16 1126–1137 Preprint at https://doi.org/10.1016/j.celrep.2016.06.059 (2016).

54. Regev, A. et al. Science forum: the human cell atlas. Elife 6, e27041 (2017).

55. Martin, C. et al. Frozen Tissue Nuclei Extraction (for 10xV3 snSEQ) v1. (2020) doi:10.17504/protocols.io.bck6iuze.

56. Balderrama, K. Nissl staining and imaging of mouse brain tissue for slide-seq registration v1. (2021) doi:10.17504/protocols.io.bv7vn9n6.

57. Stickels, R. et al. Library generation using Slide-seqV2 v1. (2020) doi:10.17504/protocols.io.bpgzmjx6.

58. 3D Slicer image computing platform. 3D Slicer https://www.slicer.org/.

59. Ungi, T., Lasso, A. & Fichtinger, G. Open-source platforms for navigated image-guided interventions. Med. Image Anal. 33, 181–186 (2016).

60. Beg, M. F., Miller, M. I., Trouvé, A. & Younes, L. Computing large deformation metric mappings via geodesic flows of diffeomorphisms. International journal of computer doi:10.1023/B:VISI.0000043755.93987.aa.

61. Tward, D., et al. Diffeomorphic Registration With Intensity Transformation and Missing Data: Application to 3D Digital Pathology of Alzheimer’s Disease. Frontiers in Neuroscience vol. 14 Preprint at https://doi.org/10.3389/fnins.2020.00052 (2020).

62. Tward, D. et al. Solving the where problem in neuroanatomy: a generative framework with learned mappings to register multimodal, incomplete data into a reference brain. bioRxiv 2020.03.22.002618 (2020) doi:10.1101/2020.03.22.002618.

63. Tward, D. J. An Optical Flow Based Left-Invariant Metric for Natural Gradient Descent in Affine Image Registration. Frontiers in Applied Mathematics and Statistics 7, (2021).

64. Zhu, D. et al. Multimodal Brain Image Analysis and Mathematical Foundations of Computational Anatomy: 4th International Workshop, MBIA 2019, and 7th International Workshop, MFCA 2019, Held in Conjunction with MICCAI 2019, Shenzhen, China, October 17, 2019, Proceedings. (Springer Nature, 2019).

65. Hahsler, M., Piekenbrock, M. & Doran, D. dbscan: Fast Density-Based Clustering with *R*. Journal of Statistical Software vol. 91 Preprint at https://doi.org/10.18637/jss.v091.i01 (2019).

66. Sparta, B., Hamilton, T., Aragones, S. D. & Deeds, E. J. Binomial models uncover biological variation during feature selection of droplet-based single-cell RNA sequencing. bioRxiv 2021.07.11.451989 (2021) doi:10.1101/2021.07.11.451989.

67. Jarvis, R. A. & Patrick, E. A. Clustering Using a Similarity Measure Based on Shared Near Neighbors. IEEE Trans. Comput. C–22, 1025–1034 (1973).

68. Ertöz, L., Steinbach, M. & Kumar, V. Finding Clusters of Different Sizes, Shapes, and Densities in Noisy, High Dimensional Data. in Proceedings of the 2003 SIAM International Conference on Data Mining (SDM) 47–58 (Society for Industrial and Applied Mathematics, 2003).

69. Traag, V. A., Van Dooren, P. & Nesterov, Y. Narrow scope for resolution-limit-free community detection. Phys. Rev. E Stat. Nonlin. Soft Matter Phys. 84, 016114 (2011).

70. Gu, Z. Complex heatmap visualization. iMeta 1, e43 (2022).

71. Davis, A., Gao, R. & Navin, N. E. SCOPIT: sample size calculations for single-cell sequencing experiments. BMC Bioinformatics 20, 566 (2019).

72. Cable, D. M. et al. Robust decomposition of cell type mixtures in spatial transcriptomics. Nat. Biotechnol. 40, 517–526 (2022).

73. Stellato, B., Banjac, G., Goulart, P., Bemporad, A. & Boyd, S. OSQP: An Operator Splitting Solver for Quadratic Programs. 2018 UKACC 12th International Conference on Control (CONTROL) Preprint at https://doi.org/10.1109/control.2018.8516834 (2018).

74. Eddelbuettel, D. Seamless R and C++ Integration with Rcpp. (Springer, 2013).

75. Karczewski, K. J. et al. Systematic single-variant and gene-based association testing of thousands of phenotypes in 394,841 UK Biobank exomes. Cell Genom 2, 100168 (2022).

76. Hail Team. Hail. (2023). doi:10.5281/zenodo.7622684.

77. Tange, O. GNU Parallel - The Command-Line Power Tool.; login: The USENIX Magazine 36, 42–47 (2011).

78. Bezanson, J., Edelman, A., Karpinski, S. & Shah, V. B. Julia: A Fresh Approach to Numerical Computing. SIAM Review vol. 59 65–98 Preprint at https://doi.org/10.1137/141000671 (2017).

79. Huangfu, Q. & Hall, J. A. J. Parallelizing the dual revised simplex method. Mathematical Programming Computation vol. 10 119–142 Preprint at https://doi.org/10.1007/s12532-017-0130-5 (2018).

80. Cplex, I. I. V22. 1: User’s Manual for CPLEX. International Business Machines Corporation.

81. Chen, E. Y. et al. Enrichr: interactive and collaborative HTML5 gene list enrichment analysis tool. BMC Bioinformatics 14, 128 (2013).

82. Xie, Z. et al. Gene Set Knowledge Discovery with Enrichr. Curr Protoc 1, e90 (2021).

83. Ashburner, M. et al. Gene ontology: tool for the unification of biology. The Gene Ontology Consortium. Nat. Genet. 25, 25–29 (2000).

84. Gene Ontology Consortium. The Gene Ontology resource: enriching a GOld mine. Nucleic Acids Res. 49, D325–D334 (2021).

85. Sayols, S. rrvgo: a Bioconductor package to reduce and visualize Gene Ontology terms. Preprint at https://ssayols.github.io/rrvgo (2020).

86. Harrell, F. E., Jr, from Charles Dupont, W. C. & others., M. Hmisc: Harrell Miscellaneous. Preprint at https://CRAN.R-project.org/package=Hmisc (2021).

87. Kanton, S. et al. Organoid single-cell genomic atlas uncovers human-specific features of brain development. Nature 574, 418–422 (2019).

88. Cao, J. et al. Joint profiling of chromatin accessibility and gene expression in thousands of single cells. Science 361, 1380–1385 (2018).

89. Zimmerman, K. D., Espeland, M. A. & Langefeld, C. D. A practical solution to pseudoreplication bias in single-cell studies. Nat. Commun. 12, 738 (2021).

90. Stuart, T. et al. Comprehensive Integration of Single-Cell Data. Cell (2019) doi:10.1016/j.cell.2019.05.031.

91. van Dijk, D. et al. Recovering Gene Interactions from Single-Cell Data Using Data Diffusion. Cell 174, 716–729.e27 (2018).

92. GeneOverlap: R package for testing and visualizing gene list overlaps. (Github).

93. Korotkevich, G. et al. Fast gene set enrichment analysis. bioRxiv 060012 (2021) doi:10.1101/060012.

94. de Leeuw, C. A., Mooij, J. M., Heskes, T. & Posthuma, D. MAGMA: generalized gene-set analysis of GWAS data. PLoS Comput. Biol. 11, e1004219 (2015).

